# In-House Forecasting of Crop Latitude Adaptation through a Daylength-sensing-based Environment Adaptation Simulator (DEAS)

**DOI:** 10.1101/2020.12.09.418558

**Authors:** Leilei Qiu, Qinqin Wu, Xiaoying Wang, Gui Zhuang, Jiupan Han, Hao Wang, Zhiyun Shang, Wei Tian, Zhuo Chen, Zechuan Lin, Hang He, Jie Hu, Qiming Lv, Juansheng Ren, Jun Xu, Chen Li, Xiangfeng Wang, Yang Li, Shaohua Li, Rongyu Huang, Xu Chen, Cheng Zhang, Ming Lu, Chengzhi Liang, Peng Qin, Xi Huang, Shigui Li, Xinhao Ouyang

## Abstract

Global climate change necessitates the accelerated breeding of new crop varieties that can sustain yields in new environments. As a proxy for environmental adaptation, the selection of crops that can adapt to different latitudes is an appealing strategy. However, such selection currently involves a lengthy procedure that severely restricts the rapid breeding of varieties. Here, we aimed to combine molecular technologies with an in-house streamlined screening method to facilitate rapid selection for latitude adaptation. We established the Daylength-sensing-based Environment Adaptation Simulator (DEAS) to measure crop latitude adaptation via the transcriptional dynamics of florigen genes at different latitudes. We used different statistical approaches to demonstrate that DEAS predicts the florigen expression profiles in rice with high accuracy. Furthermore, we demonstrated the potential for application of DEAS in different crops. Incorporating DEAS into the breeding programs of conventional and underutilized crops could help meet the future needs for crop adaptation and promote sustainable agriculture.

## INTRODUCTION

Large-scale crop migration may be an effective strategy to ensure food security in response to global warming^1–6^. Future increases in temperature may open up new agriculturally suitable areas at high latitudes, and crops at middle and low latitudes will benefit from an extension of the growth season with increasing temperatures^1, 2^. Predictably, the accelerated selection of new crop varieties for latitude adaptation might be critical for yield stability in future crop migration. The adaptation of crops to new latitudes requires retuning their flowering time to maximize yield potential^7^. To date, many studies have reported that crop latitude adaptation re-establishment always requires the genetic modification of photoperiodic genes. In potato, the natural allelic variants of StCDF1, a DOF (DNA-binding with one finger) transcription factor are no longer post-translationally regulated by light, enabling them to be cultivated outside the geographical center of origin of potato^8^. In soybean, the *J* gene is the ortholog of *Arabidopsis thaliana EARLY FLOWERING 3* (*ELF3*) and its loss-of-function variants extend the vegetative phase of soybean and improve yield under short-day conditions, thereby enabling the expansion of its cultivation in tropical regions^9^. A recent study showed accumulated impairments in the *pseudo-response-regulator* gene (PRR) orthologs *Tof11* and *Tof12* function were likely to have permitted an earlier harvest and improved adaptation to the limited summer growth period at higher latitudes during soybean domestication^10^. In maize, natural variation in the regulatory regions of *ZmCCT9*, *ZmCCT10* and *ZCN8* (an ortholog of *Arabidopsis thaliana FT*) were essential for the spread of maize from tropical to temperate regions during its early domestication^11–13^. In rice, several studies have shown that *Grain number, plant height and heading date 7* (*Ghd7*), *Heading date 1* (*Hd1*), *DAYS TO HEADING 8* (*DTH8*/*Ghd8*/*LHD1*) and *Grain number, plant height and heading date 7*.*1* (*Ghd7.1*/*DTH7*/*OsPRR37*) affect the geographical distribution of rice varieties^14–20^. Recent studies have shown that both Ghd7-Hd1 and DTH8-Hd1 modules are essential to regulate the transcription of *Ehd1* and *Hd3a*^21–25^. However, different crops have evolved different photoperiodic genes for latitude adaptation, which hampers efforts to summarize a universal mechanism underlying latitude adaptation for accelerating breeding.

The photoperiodic genes (*Tof11, Tof12, gmelf3*, *ZmCCT9-10*, *Ghd7*, *Hd1*, *DTH8* and *Ghd7.1*) ultimately enhance adaptability in different crops by directly or indirectly regulating the transcription of orthologs of the florigen gene *FT*^9–13, 15–20, 23, 26, 27^. In plants, photoperiodic genes regulate the transcription of *FT* or *FT* orthologs to sense daylength^28^. Many long-day (LD) plants such as *Arabidopsis thaliana* undergo vernalization during the winter and perceive gradually increasing daylengths in the spring, which induces flowering^29^. In Arabidopsis, the gradual increase in daylength is accompanied by a gradual accumulation of transcripts of the florigen gene *FT*^30, 31^. By contrast, typical short-day (SD) plants such as rice grow during the summer, and short days induce flowering^28, 32, 33^. Rice perceives seasonal changes via a dual-gate system that recognizes a critical daylength threshold^28^. In *japonica* rice, two light-signaling pathways in response to red and blue light involve the key genes *Ghd7* and *OsGIGANTEA* (*OsGI*) to set the daylength threshold (threshold = 13.5 h). A daylength shorter than 13.5 h induces the transcription of *Early heading date 1* (*Ehd1*) and the *FT* ortholog, *Heading date 3* (*Hd3a*)^32, 33^. Previous study showed that critical daylength sensing is widespread in plants, and different plants sense different critical thresholds to induce flowering^34^. Based on these data, we reasoned that daylength sensing based on the transcription of the florigen gene could be used as a downstream trait upon which to base a general method to select crops for latitude adaptation.

Here, we report that daylength-sensing diversity regulated by different combinations of photoperiodic genes provides multiple strategies that allow rice cultivation at different latitudes and harvest times, and improves adaptation and yield potential. Furthermore, we established a Daylength-sensing based Environment Adaptation Simulator (DEAS) that couples the transcriptional dynamics of florigen genes and latitude adaptation. Our approach allows the detection of crop daylength sensing and an assessment of crop latitude adaptation based on simulating the transcriptional dynamics of florigen genes to inform crop latitude adaptation selection and accelerate breeding.

## Results

### *Hd1hd1 dth8dth8* and *hd1hd1 DTH8dth8* Are the Two Major Genotypes of *indica* Hybrid Rice for the Improvement of Latitude Adaptation in East Asia

*Indica* rice is the main rice subspecies grown in East Asia and the development of *indica* hybrid rice varieties in recent decades has increased crop yields and provided food for a growing population^35^. Using resequencing data^36^ and haplotype analysis, we first investigated allelic variation within several essential flowering-time genes (*Hd1*^26^; *DAYS TO HEADING 2*, *DTH2*^37^; *DTH7/Ghd7*.*1/OsPRR37*^15, 16^; *DTH8*/*Ghd8*/*LHD1*^18, 19, 27^; *Ghd7*; *Hd3a*^38, 39^; *RFT1*^39^; *Heading date 6*, *Hd6*^40^; *Ehd1*^41^; *Heading date 16*, *Hd16*^42^; *Heading date 17*, *Hd17/OsELF3-1*^43–46^ and *Heading date 18*, *Hd18*^47^) in sterile and restorer lines. The results showed that *Hd1, DTH7* and *DTH8* showed functional and non-functional (i.e., mutant) allelic variation in 70 restorer lines (Fig. 1a), and *Hd1*, *DTH7*, *DTH8*, *Ghd7* and *RFT1* showed functional and non-functional allelic variation in 44 sterile lines (Fig. 1a). Notably, *indica* hybrid varieties contained only non-functional *Hd1* and *DTH8* alleles (Fig. 1a). We therefore divided the *indica* hybrid varieties into four genotypes, depending on their combinations of functional or non-functional *Hd1* and *DTH8* alleles: *Hd1hd1 DTH8dth8*, *Hd1hd1 dth8dth8*, *hd1hd1 DTH8dth8*, and *hd1hd1 dth8dth8* (Fig. 1b). A hypergeometric analysis showed that only *Hd1hd1 dth8dth8* was significantly enriched in all *indica* hybrids (Fig. 1b).

**Fig. 1.**
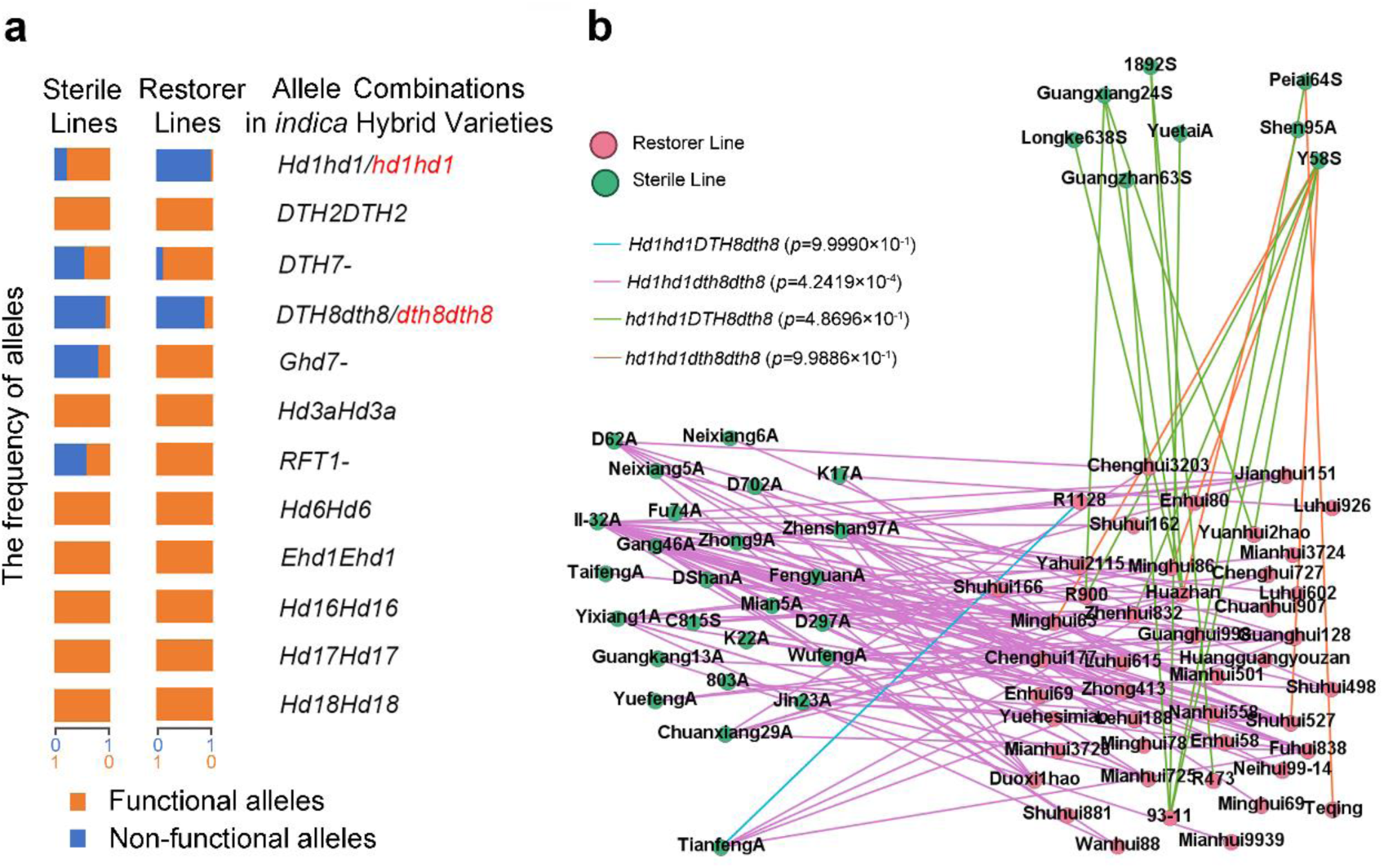
Functional and non-functional alleles of photoperiodic/flowering-time genes in *indica* hybrid rice varieties. **a** Functional and non-functional allele frequencies of flowering-time genes in sterile and restorer lines shown by hybrid allele combinations. **b** *Hd1* and *DTH8* allele combinations in *indica* hybrid rice varieties. The red hubs represent restorer lines and the green hubs represent sterile lines. Each line represents the single *indica* hybrid rice variety derived from crosses between the two parents.

Next, we assessed whether the geographic distribution of *indica* hybrid rice varieties differed, depending on the combinations of functional and non-functional *Hd1* and *DTH8* alleles in plants. Breeders sort *indica* hybrid rice distribution throughout East Asia into five ecological classes, based on geographic differences and sowing time. Class 1 describes middle-season rice cultivation in the upper reaches of the Yangtze River for one-round planting. On the basis of the critical daylength threshold (13.5 h), Class 1 rice experiences increasing and then decreasing daylengths (i.e., from SD to LD and then to SD) before flowering (Supplementary Fig. 1a–b). The mean flowering time of *indica* hybrid rice varieties in Class 1 was 112.4 days (Supplementary Fig. 2a). The hypergeometric analysis determined that hybrid varieties containing *Hd1hd1 dth8dth8* were more abundant in Class 1 than other genotypes (Supplementary Fig. 3). Class 2 designates middle-season rice cultivation in the middle and lower reaches of the Yangtze River for one-round planting. Rice planted in this area experiences daylengths that change from LD to SD before flowering (Supplementary Fig. 1c–d). The mean flowering time of *indica* hybrid rice varieties in Class 2 was 100.5 days (Supplementary Fig. 2a). Hybrid varieties with *hd1hd1 DTH8dth8* genotypes were more enriched in Class 2 (Supplementary Fig. 3). Class 3 denotes late-season rice cultivation in the middle and lower reaches of the Yangtze River for a shorter growing season. Rice planted in this area experiences decreasing daylength (from LD to SD) before flowering (Supplementary Fig. 1e–f). The mean flowering time of *indica* hybrid rice varieties in Class 3 was 83 days (Supplementary Fig. 2a). Varieties with a *hd1hd1 dth8dth8* genotype were more frequently planted in Class 3 cultivation (Supplementary Fig. 3). Class 4 designates late-season rice cultivation in the South-China subtropical zone for two-round planting. In this area, plants grow entirely in SD conditions, in which daylengths gradually shorten (Supplementary Fig. 1g–h). The mean flowering time of *indica* hybrid rice varieties in Class 4 was 84.2 days (Supplementary Fig. 2a). Class 5 indicates early-season rice cultivation in the South-China subtropical zone for two-round planting. Plants in this region grow entirely in SD conditions, in which the daylengths gradually increase (Supplementary Fig. 1i–j). The mean flowering time of *indica* hybrid rice varieties in Class 5 was 86.7 days (Supplementary Fig. 2a). We identified no significant enrichment of any genotypes in ecological Classes 4 and 5 (Supplementary Fig. 3). Intriguingly, the mean flowering time of *indica* hybrid rice varieties in ecological Classes 1 and 2 for one-round planting was longer than that in Classes 3, 4 and 5 for short-season planting or two-round planting (Supplementary Fig. 2a). Moreover, delayed flowering time in Classes 1 and 2 clearly improved the yields from one-round planting (Supplementary Fig. 2b).

We observed that the number of varieties with the *Hd1hd1 dth8dth8* genotype was greater in two combined ecological classes than other genotypes (Supplementary Fig. 4). Next, we analyzed the relationships between genotypes and cultivars suitable for cultivation in three, four, and five combined ecological classes. Remarkably, the number of hybrid rice varieties with the *Hd1hd1 dth8dth8* genotype was greater in all ecological class combinations than other genotypes (Fig. 2 and Supplementary Figs. 4–8). However, the hypergeometric analysis showed that the *Hd1hd1 dth8dth8* genotype was not significantly enriched in any ecological class combination (Supplementary Figs. 4–8). Compared with rice varieties containing *Hd1hd1 dth8dth8*, the number of hybrid varieties with the *hd1hd1 DTH8dth8* genotype was samller (Fig. 2 and Supplementary Figs. 4–8) and these varieties were more frequently enriched in all combined ecological classes (Supplementary Figs. 4–8). These results demonstrate that *Hd1hd1 dth8dth8* and *hd1hd1 DTH8dth8* are the two major genotypes of *indica* hybrid rice that can be used to improve latitude adaptation in East Asia (Fig. 2 and Supplementary Figs. 4–8).

**Fig. 2.**
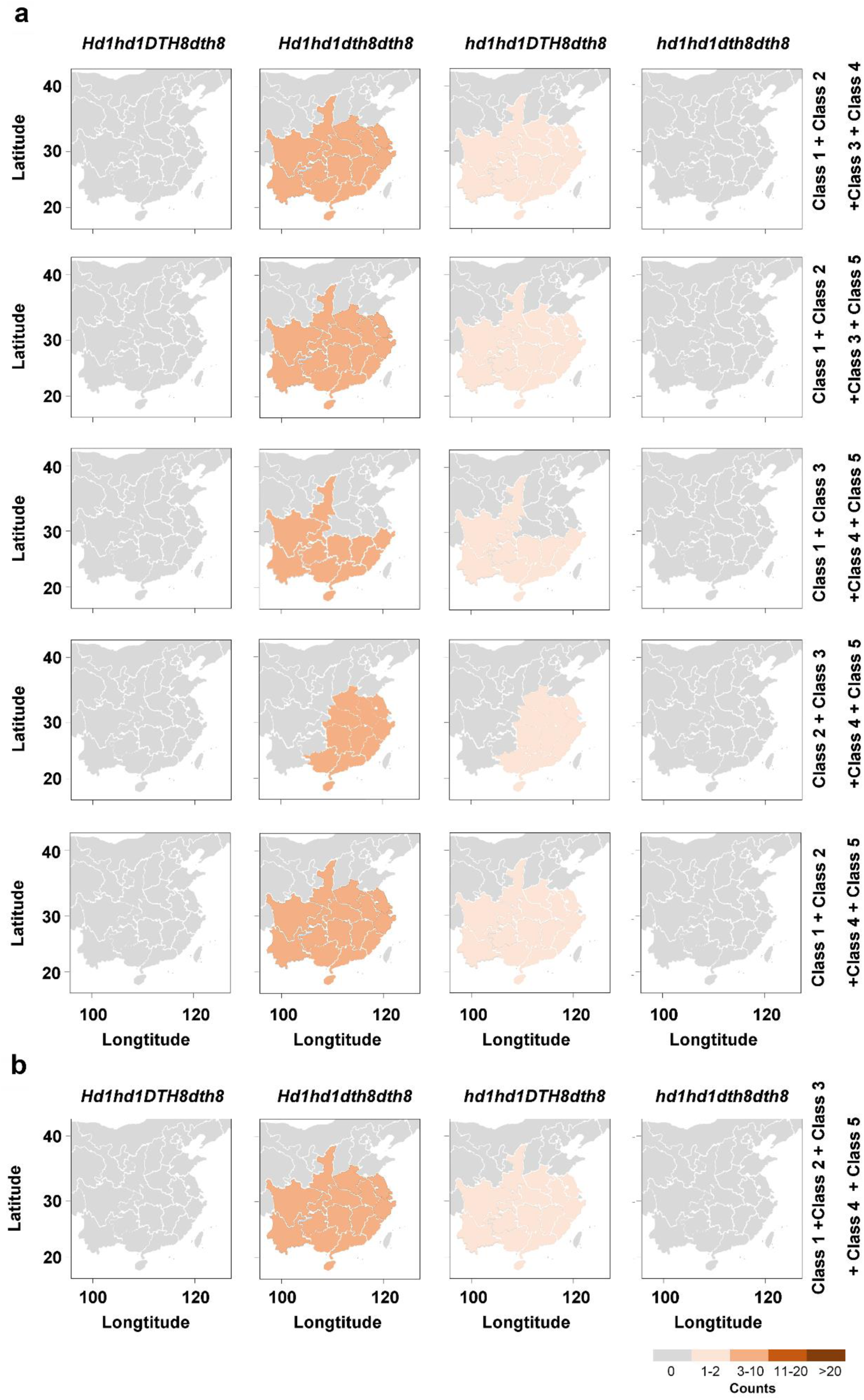
Number of *Hd1* and *DTH8* allelic combinations in *indica* hybrid rice varieties within combined ecological classes. **a** A heatmap representing the number of *Hd1* and *DTH8* allelic combinations in *indica* hybrid rice varieties in four combined ecological classes. **b** A heatmap representing the number of *Hd1* and *DTH8* allelic combinations in *indica* hybrid rice varieties in five combined ecological classes. All maps were drawn using ArcGIS 10.3 software and modified using Photoshop CC2017.

### DTH8-Hd1-Ghd7-DTH7 Represents a Core Module that Mediates Multiple Daylength-Sensing Processes in Rice

In addition to the above examples (Fig. 2), previous studies showed that non-functional *Ghd7* could enhance high-latitude adaptation in rice^15–17, 20^. Furthermore, *Ghd7* can set the critical daylength threshold in rice^32, 33^ and can regulate flowering time by interacting with Hd1 and DTH8^21–25, 48, 49^. Thus, we first investigated the genetic relationship between Ghd7 and the DTH8-Hd1 module. The results showed that *Ghd7* enhanced the repressive activity of the DTH8-Hd1 module on flowering in LD (Supplementary Fig. 9a–b). Yeast two-hybrid and luciferase complementation imaging (LCI) assays showed that Ghd7, Hd1, and DTH8 can interact with each other (Supplementary Fig. 9c–d). Therefore, together with previous studies^21–25, 48, 49^, the results support the conclusion that the DTH8-Hd1-Ghd7 module represses flowering in LD.

We also investigated whether the DTH8-Hd1-Ghd7 module affects daylength sensing. To do this in a controlled manner, we generated a Daylength-sensing based Environment Adaptation Simulator (DEAS) to simulate increasing or decreasing daylength conditions (Fig. 3a and 4a, Methods). In DEAS Step1 (Fig. 3a), we transferred plants from SD to LD conditions (to simulate increasing daylength) (Fig. 3a). Conversely, to simulate decreasing daylength, we transferred plants from LD to SD conditions (Fig. 3a). To gain insight into which mechanisms underlie daylength perception, we measured whether transcript levels of *FT* orthologs (*Hd3a,* and *RICE FLOWERING LOCUS T 1* (*RFT1*)) changed in response to daylength.

**Fig. 3.**
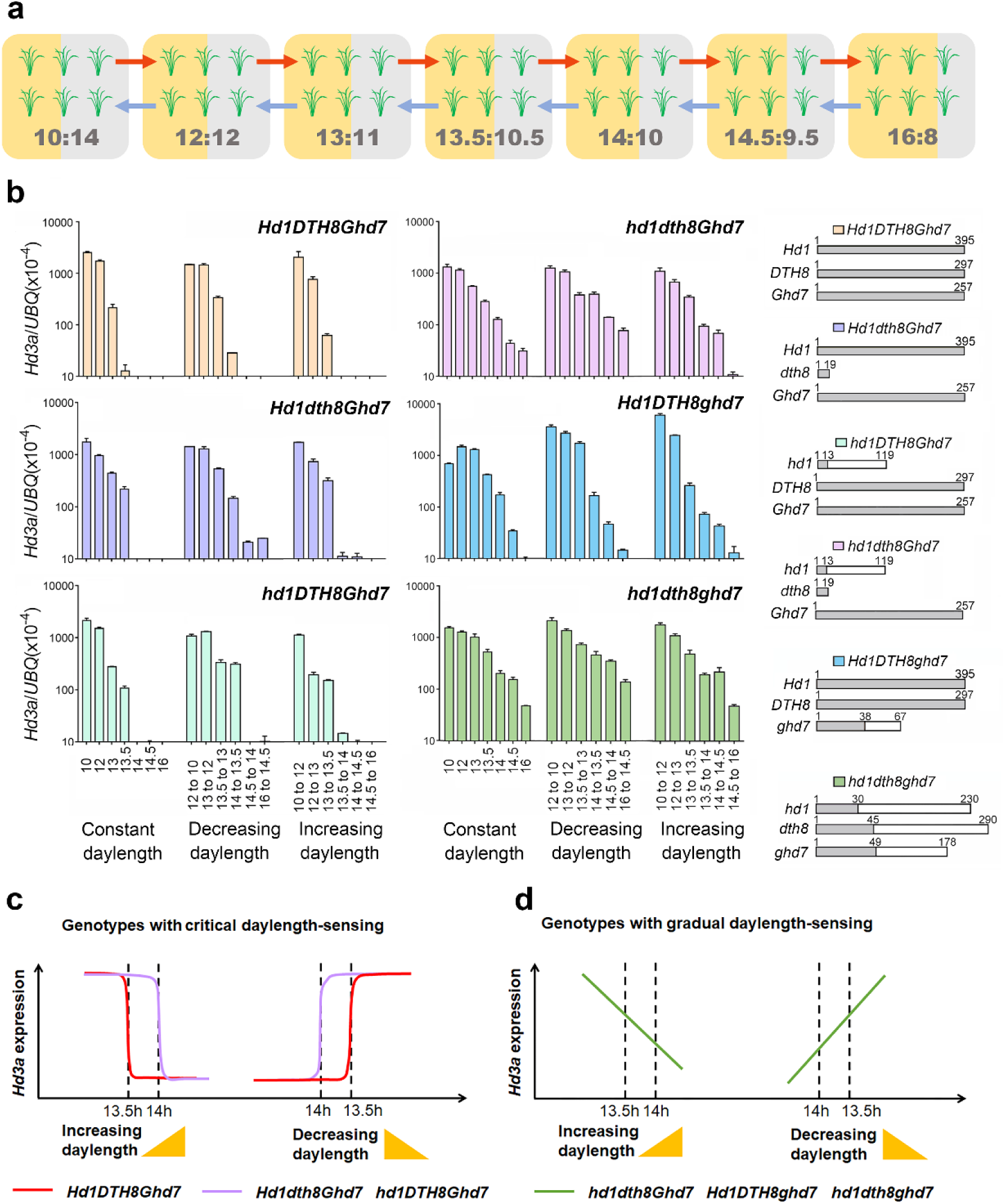
Using DEAS Step1 to detect rice daylength-sensing processes. **a** Diagram of DEAS Step1: after rice seedlings were grown for 28 days, subgroups were transferred to different growth chambers to simulate changes in daylength. Red and blue arrows indicate increasing or decreasing daylengths, respectively. **b** *Hd3a* levels under various daylength conditions. Relative mRNA levels were determined by RT-qPCR. Error bars represent the SD of three biological replicates. Gray rectangles show the lengths of the Hd1, DTH8 or Ghd7 proteins in numbers of amino acids, respectively. White rectangles represent proteins resulting from frameshift mutations in *Hd1* or *DTH8*. Small gray rectangles represent a proteins resulting from a premature stop codon in *Hd1* or *DTH8*. DNA sequencing results are shown in Supplementary Fig. 20. **c–d** Schematic diagram of different daylength-sensing processes in different genotypes.

**Fig. 4.**
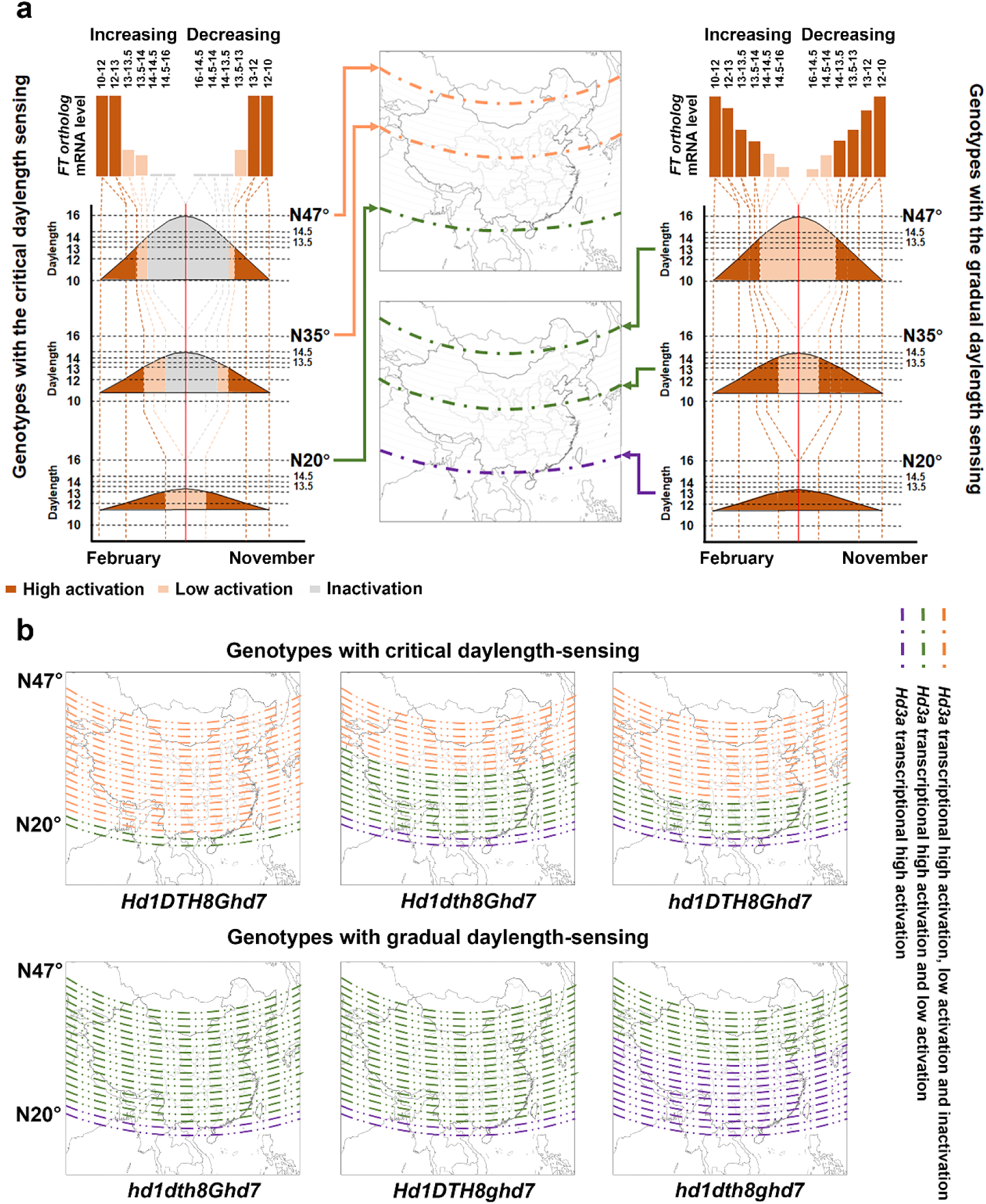
DEAS Step2 couples environmental adaptation and daylength sensing. **a** Diagram of DEAS Step2: The *FT* ortholog expression dynamic was first generated using DEAS Step1 data, then the adaptability of rice at different latitudes was described using the dynamics of *FT* ortholog expression. **b** Maps illustrating different *Hd3a* expression dynamics at different latitudes for different genotypes. All maps were drawn using ArcGIS 10.3 software and modified using Photoshop CC2017.

Daylength lengthening gradually induced *Ghd7* transcription (Supplementary Fig. 10a–c), which was similar to the findings of previous studies^32, 33^. In the *Hd1 DTH8 Ghd7* wild type (Dongjin background), constant daylengths shorter than 13.5 h induced *Ehd1*, *Hd3a*, and *RFT1* transcription (Fig. 3b and Supplementary Fig. 10d and g). Increasing and decreasing daylengths (i.e., transferring plants from 10 to 12 h, 12 to 13 h, 13.5 to 13 h, 13 to 12 h, and 12 to 10 h) induced *Ehd1*, *Hd3a*, and *RFT1* transcription (Fig. 3b and Supplementary Fig. 10e and h). However, increasing and decreasing daylengths that remained longer than the critical threshold (i.e., when plants were transferred from 13 to 13.5 h, 13.5 to 14 h, 14 to 14.5 h, 14.5 to 16 h, 14.5 to 14 h and 16 to 14.5 h) did not induce transcription (Fig. 3b and Supplementary Fig. 10e, f, h and i). Seedlings with an *Hd1 DTH8 Ghd7* genotype sensed a critical daylength threshold and induced expression of *Hd3a* and *RFT1* only if the daylength was shorter than 13.5 h (Fig. 3b–c). Hereafter, we refer to wild-type (*Hd1 DTH8 Ghd7* genotype) responses as original critical daylength sensing.

In the *Hd1 dth8 Ghd7* and *hd1 DTH8 Ghd7* genotypes, the induction of *Ehd1* transcription did not depend on the critical daylength threshold (Supplementary Fig. 10d–f); even though daylengths were longer than 13.5 h, plants maintained high levels of *Ehd1* expression (Supplementary Fig. 10d–f). However, daylengths shorter than 14 h induced *Hd3a* and *RFT1* transcription in *Hd1 dth8 Ghd7* and *hd1 DTH8 Ghd7* (Fig. 3b–c and Supplementary Fig. 10h). In particular, transferring plants from 14 to 13.5 h increased *Hd3a* and *RFT1* expression in *Hd1 dth8 Ghd7* and *hd1 DTH8 Ghd7* (more than the same transfer affected the *Hd1 DTH8 Ghd7* genotype; Fig. 3b–c and Supplementary Fig. 10h), indicating that the critical daylength threshold in *Hd1 dth8 Ghd7* and *hd1 DTH8 Ghd7* had increased to 14 h (Fig. 3c). Thus, seedlings with the *Hd1 dth8 Ghd7* and *hd1 DTH8 Ghd7* genotypes sensed a fine-tuned critical daylength threshold and induced expression of *Hd3a* and *RFT1* only if the daylength was shorter than 14 h (Fig. 3b–c).

In the *hd1 dth8 Ghd7* double mutant, *Ehd1*, *Hd3a*, and *RFT1* transcription did not depend on any critical daylength threshold (Fig. 3b, d and Supplementary Fig. 10d–i). Even at daylengths longer than 13.5 h or 14 h, these plants maintained high levels of *Ehd1*, *Hd3a*, and *RFT1* expression (Fig. 3b, d and Supplementary Fig. 10d–i); however, *hd1 dth8 Ghd7* still perceived gradual changes in daylength (Fig. 3b, d). When daylengths gradually decreased, *Hd3a* expression increased accordingly (Fig. 3d). Thus, seedlings with a *hd1 dth8 Ghd7* genotype sensed the gradual changes in daylength and induced expression of *Hd3a* and *RFT1* when daylengths gradually decreased (Fig. 3d and Supplementary Fig. 10).

In the *Hd1 DTH8 ghd7* and *hd1 dth8 ghd7* genotypes, the induction of *Ehd1* transcription did not depend on any critical daylength threshold (Supplementary Fig. 10d–f); even though daylengths were longer than 13.5 h or 14 h, plants maintained high levels of *Ehd1* expression (Supplementary Fig. 10d–f). In the presence of active Ghd7, the original critical daylength threshold was perceived in *Hd1 DTH8 Ghd7* (Fig. 3c); however, the expression of *Hd3a* and *RFT1* became insensitive to any critical daylength threshold in *Hd1 DTH8 ghd7* and *hd1 dth8 ghd7*, although it remained sensitive to gradual changes in daylength (Fig. 3d and Supplementary Fig. 10g–i). These results indicated that the DTH8-Hd1 module depends on Ghd7 to sense the critical daylength threshold.

Previous studies also showed that non-functional *DTH7* could enhance rice high-latitude adaptation^15–17^ and that DTH7 can interact with DTH8 and NF-YC1/7^25^. The cultivar Hoshinoyume (HOS), which possesses non-functional *Ghd7* and *DTH7* alleles, exhibited daylength insensitivity to promote *Hd3a* expression continuously in different daylength treatments^33^. Moreover, the nearly isogenic line NIL (*Ghd7*), which carries a functional *Ghd7* allele, mimicked a *dth7* single mutant that exhibited gradual daylength sensing^33^. These results indicated that *DTH7* regulates daylength sensing in rice. A recent study^36^ and our results showed that 93-11 carried a non-functional *DTH7* allele (Fig. 1b). Using DEAS step1, we detected the daylength-sensing processes in NIL*^Hd1 DTH8 Ghd7^* (mimics the *dth7* single mutant), NIL*^Hd1 dth8 Ghd7^* (mimics *dth8 dth7*), NIL*^hd1 DTH8 Ghd7^* (mimics *hd1 dth7*) and NIL*^hd1 dth8 Ghd7^* (mimics *hd1 dth8 dth7*). The results showed that NIL*^Hd1 DTH8 Ghd7^* (mimics *dth7*) exhibited critical daylength sensing (threshold = 12 h) of *RFT1* expression and gradual daylength sensing of *Hd3a* expression (Supplementary Fig. 11). NIL*^hd1 DTH8 Ghd7^* (mimics *hd1 dth7*) exhibited gradual daylength sensing of *RFT1* expression and daylength insensitivity with high *Hd3a* mRNA levels (Supplementary Fig. 11). NIL*^Hd1 dth8 Ghd7^* (mimics *dth8 dth7*) and NIL*^hd1 dth8 Ghd7^* (mimics *hd1 dth8 dth7*) exhibited daylength insensitivity with high *Ehd1, Hd3a* and *RFT1* mRNA levels (Supplementary Fig. 11). Therefore, the data confirm that the DTH8-Hd1-Ghd7-DTH7 is a core module that mediates multiple daylength-sensing processes in rice.

### DEAS Couples Latitude Adaptation and Daylength Sensing

To investigate the relationship between latitude adaptation and daylength sensing, we collected daylength data for the whole year between N47° and N20°. Because the rate of change in daylength is not constant, but rather is fastest in spring and autumn (Supplementary Fig. 12a), the degree of precision varies with the time of year. The rate of change is lower at low latitudes during much of the year and is higher at high latitudes (Supplementary Fig. 12a). This illustrates the one-to-one correlation between latitude and the changing daylength profile. We demonstrated that expression of the *FT* ortholog (*Hd3a* or *RFT1*) responds to changing daylength (Fig. 3 and Supplementary Figs. 10 and 11). On this basis, we designed DEAS Step2, which can infer the dynamic of *Hd3a* (or *RFT1*) expression that corresponds to natural daylength changes at one latitude with respect to expression of the *FT* ortholog (*Hd3a* or *RFT1*) in increasing and decreasing daylengths in DEAS Step1 (Fig. 3a). The dynamics of *Hd3a* (or *RFT1*) expression at different latitudes can be used to accurately forecast the latitude adaptability of rice with different daylength-sensing processes on the map using DEAS Step2 (Fig. 4a, Methods).

As the latitude increases, the duration of the photoperiod in summer is longer (Supplementary Fig. 12a). Therefore, rice cultivars adapted to middle and high latitudes delay flowering to make full use of the longer duration of LD for maximal productivity. According to DEAS, although the dynamics of *Hd3a* expression in rice of any genotype showed a longer duration of inhibition or lower activation with the increase in latitude (Fig. 4a, b and Supplementary Fig. 12b), we observed that rice with critical daylength sensing exhibited stronger transcriptional repression of *FT* orthologs at middle and high latitudes compared to those that used gradual daylength sensing (Fig. 4a). Notably, transcriptional inactivation of *Hd3a* and *RFT1* was observed only for *Hd1 DTH8 Ghd7*, *hd1 DTH8 Ghd7* and *Hd1 dth8 Ghd7* via critical daylength sensing (Fig. 4b and Supplementary Figs. 13–17). These results indicated that rice with critical daylength sensing can adapt to middle and high latitudes by delaying flowering and improving yield potential.

Conversely, rice with gradual daylength sensing exhibited a stronger transcriptional activation of *FT* orthologs compared with rice that used critical daylength sensing (Fig. 4a). Even at middle and high latitudes with long photoperiods, the genotypes that possessed gradual daylength sensing still activated *FT* ortholog expression to promote flowering (Fig. 4a). In *hd1 dth8 Ghd7*, *Hd1 DTH8 ghd7* and *hd1 dth8 ghd7* genotypes that displayed gradual daylength sensing, the dynamics of *Hd3a* and *RFT1* expression mainly exhibited high and low activation or constantly high activation at different latitudes (Fig. 4b and Supplementary Figs. 13–17). Therefore, genotypes that possess gradual daylength sensing result in more rapid growth cycles that allow adaptation to the shorter growing seasons at high latitude, or multiple plantings in longer seasons at middle and low latitudes.

Because *Hd1 DTH8 Ghd7* genotypes sensed the original critical daylength threshold (13.5 h), the dynamics of *Hd3a* expression still exhibited low activation or inactivation at a low latitude (N23°) (Fig. 4b and Supplementary Fig. 13). Thus, delayed flowering caused by the low expression of *Hd3a* in *Hd1 DTH8 Ghd7* would not allow adaptation to multiple plantings in longer seasons at low latitudes. However, *Hd1 dth8 Ghd7* and *hd1 DTH8 Ghd7*, which sensed a 14-h threshold, repressed *Hd3a* expression during the LD period at high and mid-latitudes (N47° to N30°30′) (Fig. 4b and Supplementary Fig. 13). As the latitude decreased, a gradual increase in the activation of *Hd3a* at low latitudes was observed in genotypes that sense a 14-h threshold (Fig. 4b and Supplementary Fig. 13). Similar dynamics of *RFT1* expression at different latitudes are shown in Supplementary Fig. 17. Consequently, the promotion of flowering time caused by high expression of *Hd3a* and *RFT1* in *Hd1 dth8 Ghd7* and *hd1 DTH8 Ghd7* could allow multiple plantings in longer seasons at low latitudes. The fine-tuning of the critical daylength threshold might enable rice to more precisely regulate *Hd3a* expression with regard to latitude and improve adaptability.

To explain why *Hd1hd1 dth8dth8* and *hd1hd1 DTH8dth8* genotypes can be cultivated in multiple ecological classes, we analyzed the relationship between the *Hd3a* expression pattern and latitude in *Hd1hd1 dth8dth8* and *hd1hd1 DTH8dth8* genotypes using DEAS Step2 inference. Rice seedlings with *Hd1hd1 dth8dth8* and *hd1hd1 DTH8dth8* genotypes sensed a critical daylength threshold and induced expression of *Hd3a* only if the daylength was shorter than 13 h (Supplementary Fig. 18a). We observed that the *Hd1hd1 dth8dth8* and *hd1hd1 DTH8dth8* genotypes exhibited different dynamics of *Hd3a* transcription in response to ecological classes (Supplementary Fig. 18b), which was similar to those of *Hd1 dth8 Ghd7* and *hd1 DTH8 Ghd7* (Fig. 4b and Supplementary Fig. 13–17). This enabled them to flower more efficiently at different latitudes and improved their adaptation and yield potential to different environments.

### Evaluation of DEAS Inference Accuracy

The most crucial feature of DEAS is that it can be used together with gene expression data to infer the dynamics of *FT* ortholog expression (Fig. 4a). To assess the accuracy of DEAS inference, we individually simulated the daylength change dynamics of five ecological classes in growth chambers (Fig. 5). Based on Fig. 4a, we further inferred the dynamics of *Hd3a* expression in response to the five photoperiodic regimes of the five ecological classes in different genotypes (Fig. 5). Next, the time-course of *Hd3a* expression was detected in each of the simulated five ecological classes (Fig. 5). To satisfy different requirements of breeders and scientific researchers, we used two different methods to evaluate the accuracy of DEAS prediction: the categorical manner and the continuous manner (Methods).

**Fig. 5.**
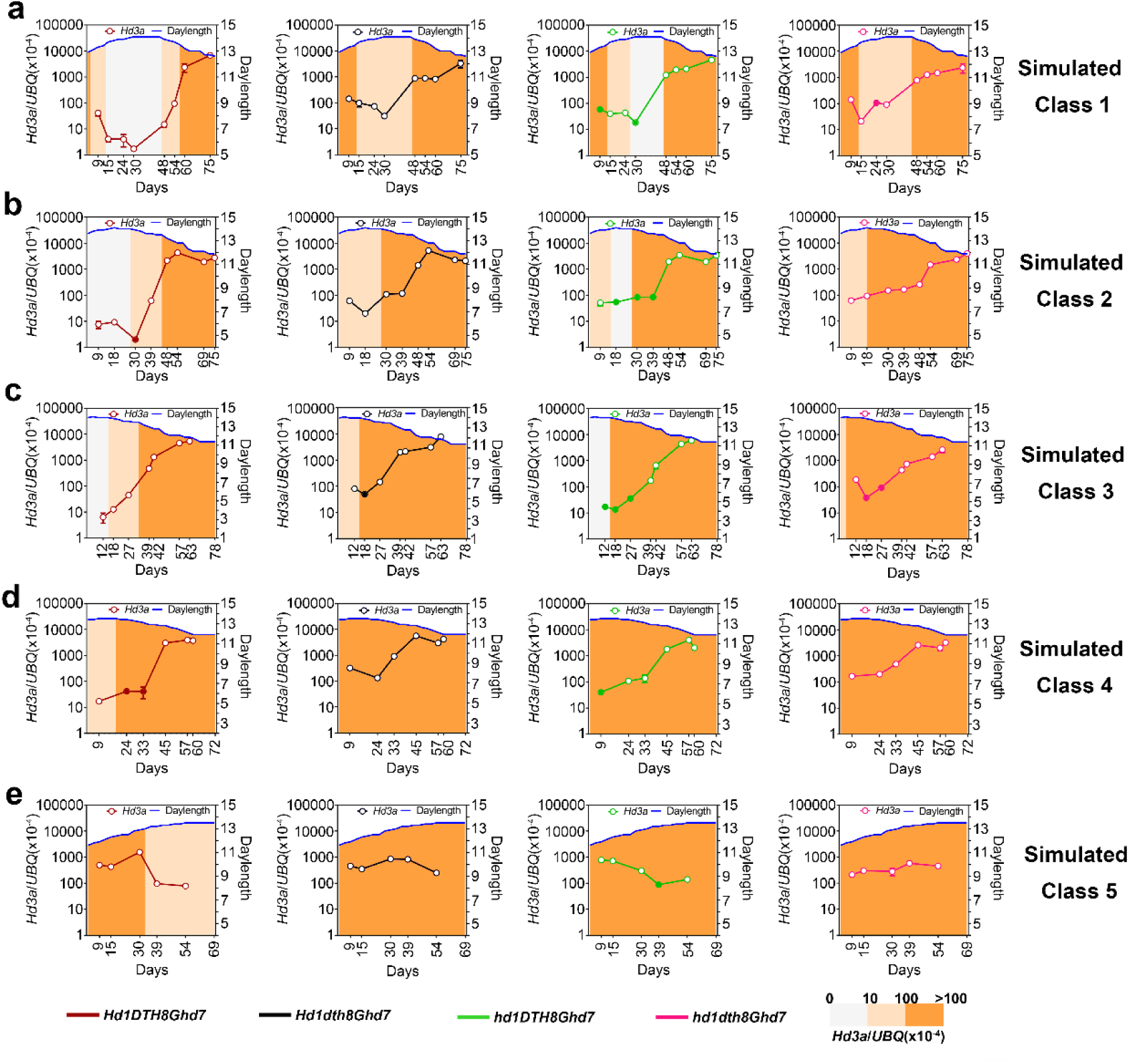
The categorical manner for evaluating DEAS inference accuracy using *Hd3a* mRNA levels in simulated five ecological classes. The relative mRNA level determined by RT-qPCR. Error bars represent the SD for three biological replicates. Blue lines represent changing daylength in the simulated five ecological classes. The four different colored lines represent the estimated values using DEAS. The filled and empty circles indicate that the data were consistent or inconsistent, respectively, with the inference represented by the three background colors.

Using the categorical manner, the real *Hd3a* and *RFT1* expression data and DEAS Step2 inference exhibited 87.5% and 89.7% coincidence, respectively (Fig. 5 and Supplementary Figs. 19). Using the continuous manner, the coefficient of determination *r^2^* showed that DEAS Step2 could accurately infer the *Hd3a* and *RFT1* expression profiles in simulated Classes 1–3 (Supplementary Figs. 20–21). However, the continuous manner failed to infer the *Hd3a* and *RFT1* expression profiles well in simulated Classes 4–5 (Supplementary Figs. 20–21). Thus, these results indicated that the DEAS was more accurate in the quantitative inference of mid-high latitude with large variations in daylength. On the contrary, at low latitudes with small variations in daylength (<13.5 h), continuous high *Hd3a* and *RFT1* expression led to the failure of the quantitative prediction of the model.

The cost of building and running seven temperature-controlled growth chambers is very high (Fig. 3a). The DEAS could be set up and applied in breeding institutes at a lower cost, and all seedlings could be grown at a constant temperature in DEAS step1 in all simulated classes (Methods). However, because the temperature is highly variable under real field conditions, it needs to be determined whether the DEAS could infer *Hd3a* and *RFT1* expression profiles in the field independently from other factors such as temperature. Based on previously published data for a rice paddy field at Tsukuba, Japan grown under normal agricultural conditions^50^, the DAES inference of the *Hd3a* and *RFT1* expression profiles was more accurate (*p*<0.01) according to the continuous manner (Fig. 6a–d). However, the DEAS did not fully fit all data (*r*^2^*_Hd3a_*=0.788 and *r*^2^*_RFT1_*=0.534). We conclude that temperature and/or other factors weakly interfered with the accuracy of field data prediction in DEAS. Furthermore, we used Fisher’s Exact Test to compare the inference accuracy between simulated classes (Fig. 5 and Supplementary Figs. 19) and in the paddy field in the categorical manner. Although temperature and other factors play a role in paddy field conditions, there were respectively 82.97% (*p_Hd3a_*=0.8297) and 82.87% (*p_RFT1_*=0.8287) probabilities that the DEAS inference was consistent with that of the simulated classes and the paddy field based on Fisher’s Exact Test (Fig. 6e–f). These results confirm that daylength sensing is still the most important factor with which to infer the florigen expression profile for crop latitude adaptation.

**Fig. 6.**
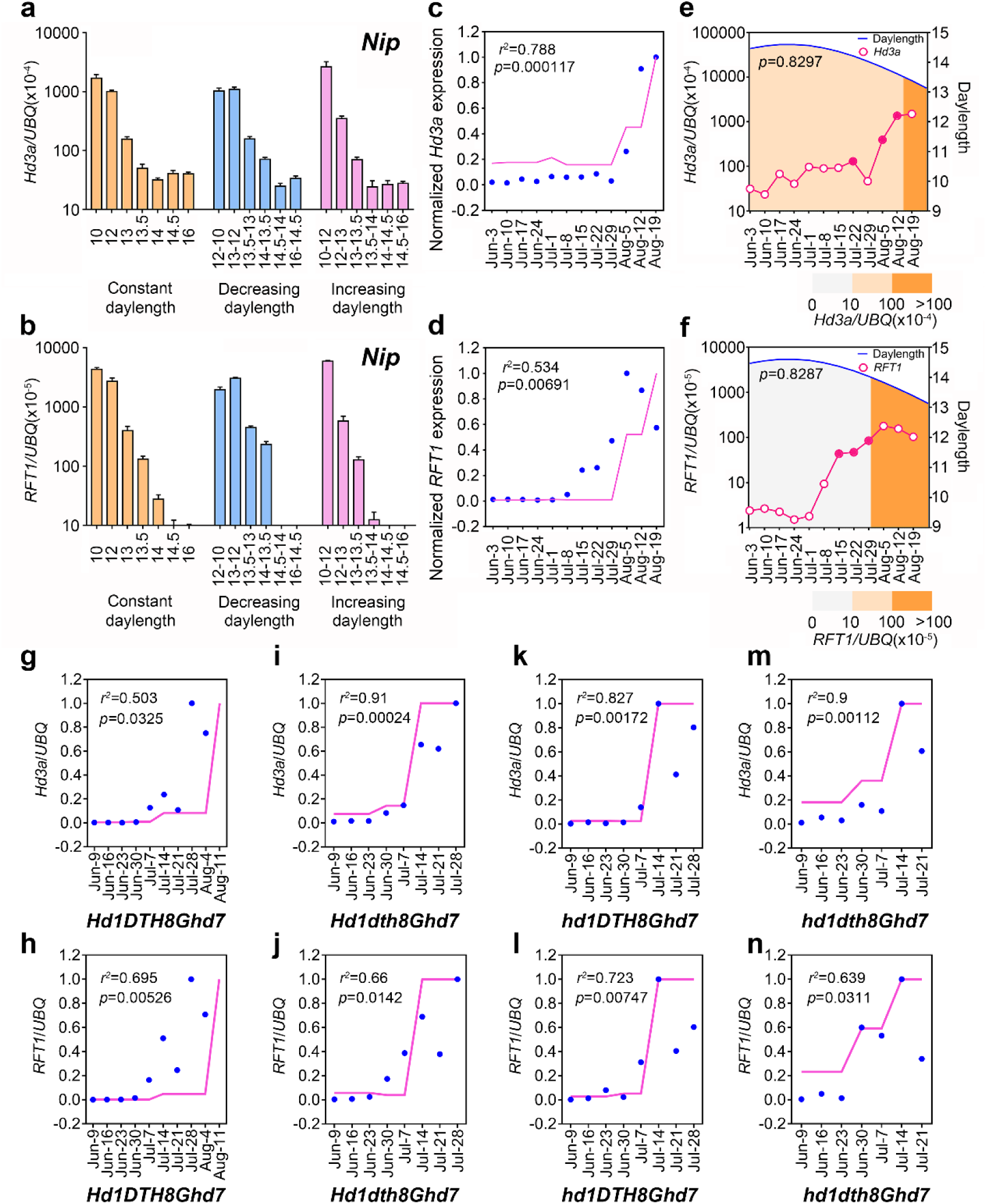
Evaluation of DEAS inference accuracy using field data. **a–b** *Hd3a* and *RFT1* mRNA levels in Nipponbare under various daylength conditions in DEAS Step1. Relative mRNA levels were determined by RT-qPCR. Error bars represent the SD for three biological replicates. **c–d** The continuous manner for evaluating DEAS inference accuracy using field data at Tsukuba, Japan. Blue points represent experimentally measured *Hd3a* mRNA levels in five simulated ecological classes. Pink lines represent the estimated value using DEAS. The coefficient of determination *r*^2^ and *p*-value can be calculated to evaluate DEAS inference accuracy **e–f** The categorical manner for evaluating DEAS inference accuracy using field data. at Tsukuba, Japan. Blue lines represent changing daylength in field condition. Pink lines represent the estimated value using DEAS. Pink filled circles and hollow circles indicate that field data were consistent or inconsistent, respectively, with the inference represented by three background colors. **g-n** The continuous manner for evaluating DEAS inference accuracy using field data at Sichuan, China. Blue points represent experimentally measured *Hd3a* mRNA levels in five simulated ecological classes. Pink lines represent the estimated value using DEAS. The coefficient of determination *r*^2^ and *p*-value can be calculated to evaluate DEAS inference accuracy

Furthermore, based on data for a rice paddy field at Sichuan, China (Class 1), the DAES inference for the *Hd3a* and *RFT1* expression profiles was more accurate (all *r*^2^ >0.05, all *p*<0.05) according to the continuous manner (Fig. 6g–n). Our results also showed that the inference accuracy (all *r*^2^ >0.05, all *p*<0.05) of the DEAS was similar between *Hd1 DTH8 Ghd7* (*r*^2^*_Hd3a_*=0.503 and *r*^2^*_RFT1_*=0.695), *Hd1 dth8 Ghd7* (*r*^2^*_Hd3a_*=0.91 and *r*^2^*_RFT1_*=0.66) and *hd1 DTH8 Ghd7* (*r*^2^*_Hd3a_*=0.827 and *r*^2^ =0.723) with critical daylength sensing (Fig. 6g-l) and *hd1 dth8 Ghd7* (*r*^2^*_Hd3a_*=0.9 and *r*^2^*_RFT1_*=0.639) with gradual daylength sensing (Fig. 6m-n). In addition, the results demonstrated that the DEAS can accurately predict the latitude adaptation of different genotypes with different daylength sensing processes.

### Potential Application of DEAS to Other Crops

We explored the potential application of DEAS to other crops. Firstly, we identified the daylength-sensing mechanism of two soybean varieties Taiwan 1 and Heilongjiang 64. In DEAS step1, the expression pattern of one soybean *FT* ortholog, *GmFT2a*, exhibited critical daylength sensing (14.5-h threshold) in Taiwan 1 (Fig. 7a). By contrast, Heilongjiang 64 was insensitive to changes in daylength at a continuously high expression level of *GmFT2a* (Fig. 7a). Next, DEAS step2 showed that the two soybean varieties exhibited different dynamics of *GmFT2a* at high latitude. TaiWan1, which possessed critical daylength sensing, showed consistent repression of *GmFT2a* expression from 1 May to 1 Sep at high latitude (N47°) (Fig. 7b). On the contrary, Heilongjiang 64 showed consistently induced *GmFT2a* expression from 1 May to 1 Sep at high latitude (N47°) (Fig. 7b). This indicates that Heilongjiang 64 is suitable for planting at high latitudes (>N40°) that have cool summers and a short growth season. Furthermore, DEAS also successfully identified the daylength-sensing mechanism of two maize varieties (Supplementary Fig. 22). Therefore, these results demonstrate the potential of DEAS in elucidating the adaptability of other crops.

**Fig. 7.**
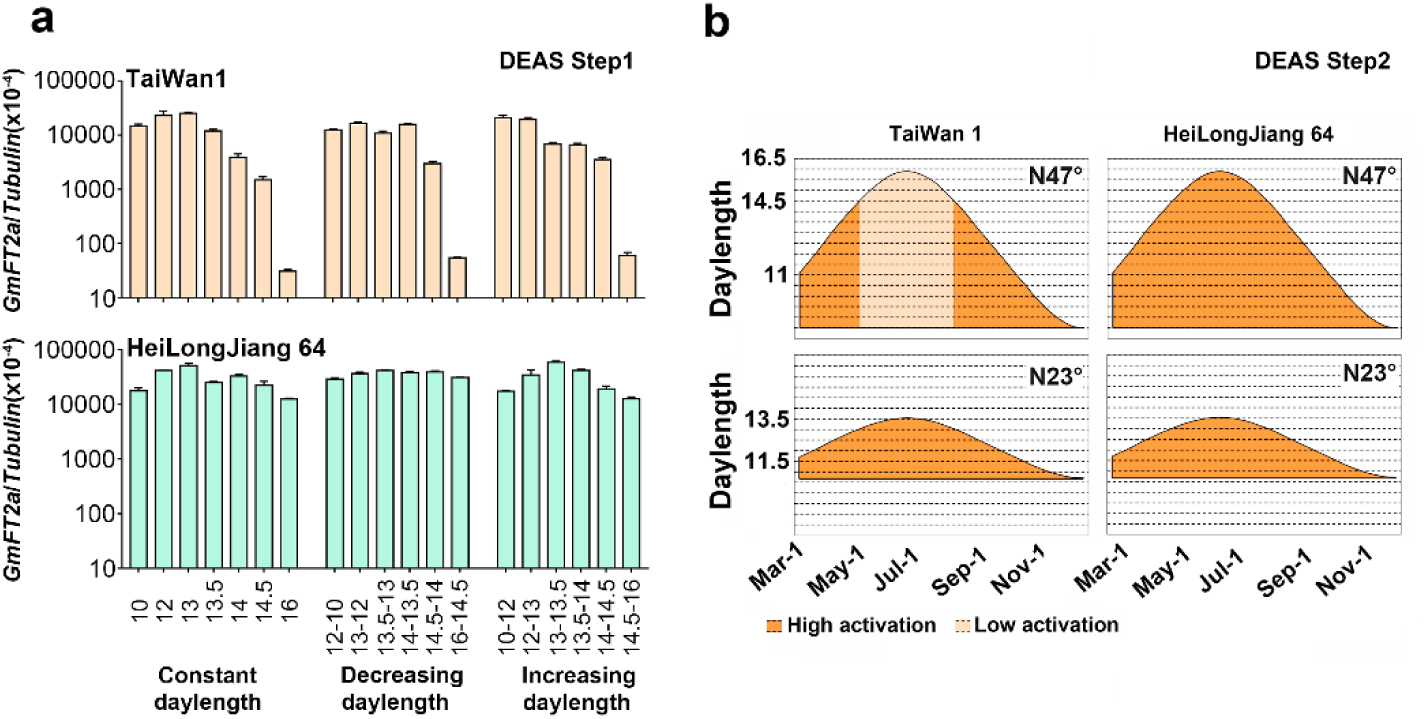
Application of DEAS to other crops. **a** *GmFT2a* mRNA levels under various daylength conditions in DEAS Step1. Relative mRNA levels were determined by RT-qPCR. Error bars represent the SD for three biological replicates. **b** Dynamics of *GmFT2a* transcription in response to different latitudes in DEAS Step2.

## Discussion

### Rice Latitude Adaptation in East Asia

Latitude adaptation refers to actions that exploit beneficial flowering time under a given reproductive cycle to maximize yield. As it is used here, adaptation refers exclusively to human activities, which include accelerating the evolutionary processes by which organisms are better fitted to their environment (e.g., selective breeding)^1^. In this study, we propose that the core photoperiodic regulators also play pivotal roles in agriculture, especially those that regulate florigen expression in response to daylength stimuli for latitude adaptation. In rice, the DTH8-Hd1-Ghd7-DTH7-mediated diversity in daylength sensing of florigen expression offers multiple strategies for the adaptability of rice to different latitudes (Fig. 8). Notably, *indica* cultivars that carry the *hd1* and *dth8* mutations can fine-tune their critical daylength threshold, which enables them to adapt to middle and high latitudes by delaying flowering and improving yield potential for one-round planting (Fig. 8a and c). However, the wild-type *indica* cultivars (*Hd1 DTH8 Ghd7 DTH7*) showed severely delayed flowering and did not head even after 24 October, when the low temperature is unfavourable for rice growing in Wuhan (N30°30′)^48^. Similarly, the wild-type *japonica* cultivar Dongjin (*Hd1 DTH8 Ghd7 DTH7*), with original critical daylength sensing, cannot adapt to high latitudes (>N53°)^17^. The late-flowering phenotype prevents the wild-type cultivars from producing seeds before winter (Fig. 8b). Indeed, the wild-type *japonica* cultivars sense the original critical daylength threshold, which ensures that they grow at mid and high latitudes (<N40°) (Fig. 8c), with a longer period of vegetative growth^17^.

**Fig. 8.**
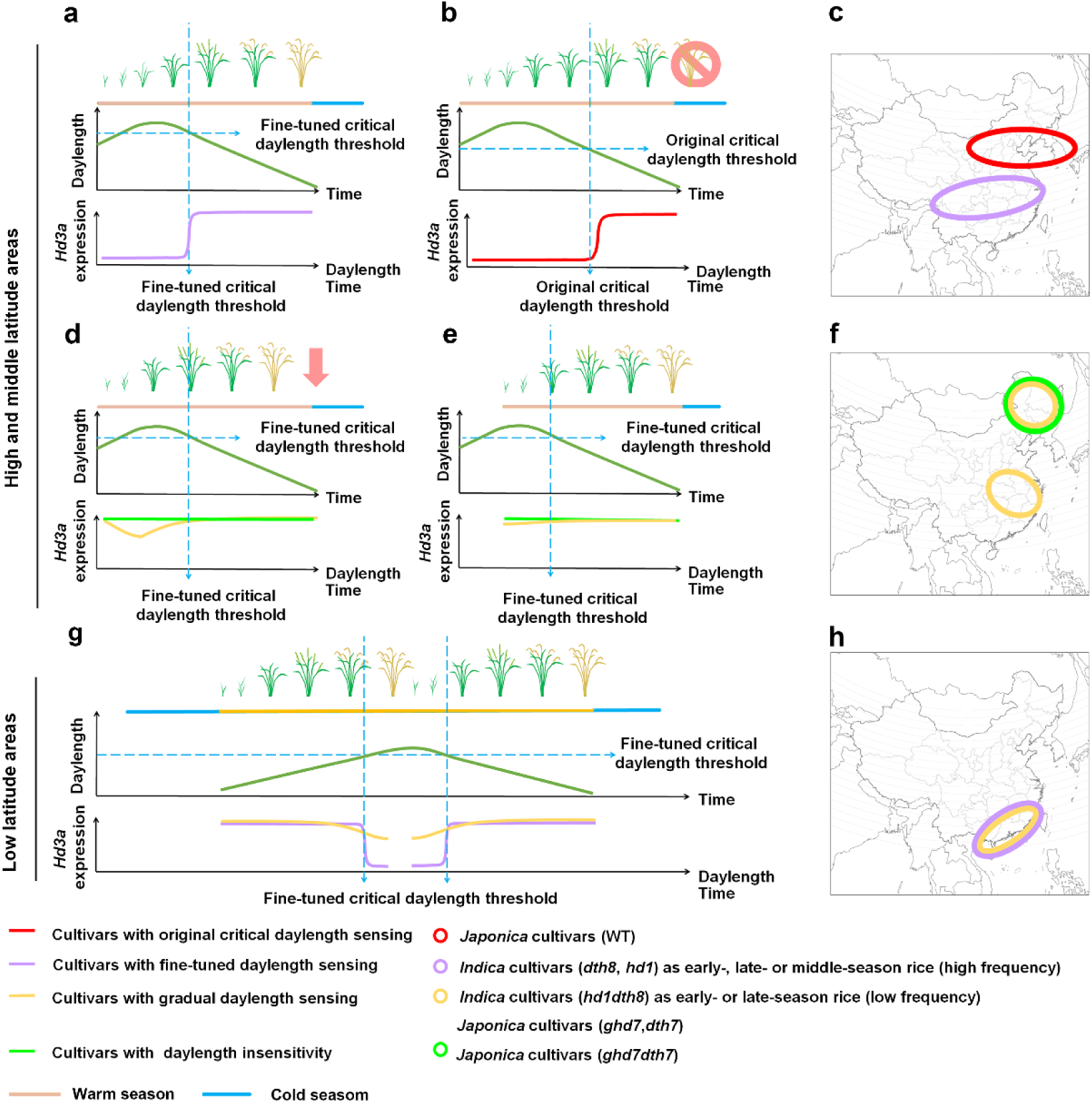
A working model to illustrate how daylength-sensing diversity regulated by photoperiod mediates rice latitude adaptation. **a–b** *hd1* (*indica*) and *dth8* (*indica*) with fine-tuned critical daylength sensing enables adaptation to middle and high latitudes by delaying flowering and improving yield potential for one-round planting. The wild type (*indica*) with critical daylength sensing shows severely delayed flowering that does not enable them to adapt to middle and high latitudes. **c** Maps illustrating the distribution of the wild type (*japonica*), *hd1* (*indica*) and *dth8* (*indica*) in East Asia. **d–e** *ghd7* (*japonica*), *dth7* (*japonica*) and *hd1 dth8* (*indica*) with gradual daylength sensing and *ghd7 dth7* (*japonica*) with daylength insensitivity undergo faster growth cycles that allow adaptation to shorter growth seasons. **f** Maps illustrating the distribution of the *ghd7* (*japonica*), *dth7* (*japonica*), *hd1 dth8* (*indica*) and *ghd7 dth7* (*japonica*) in East Asia. **g** *hd1* (*indica*) and *dth8* (*indica*) with fine-tuned critical daylength sensing and *hd1 dth8* (*indica*) with gradual daylength sensing adapt to low latitudes via the promotion of flowering and rapid life cycles for two-round planting. **h** Maps illustrating the distribution of *hd1* (*indica*), *dth8* (*indica*) and *hd1 dth8* (*indica*) in East Asia. All maps were drawn using ArcGIS 10.3 software and modified using Photoshop CC2017.

The genotypes with gradual daylength sensing can perform more rapid growth cycles that allow adaptation to shorter growing seasons (Fig. 8d–e). *Japonica* cultivars carrying *ghd7* single mutations exhibit gradual daylength sensing^33^ and those carrying *ghd7 dth7* double mutations exhibit daylength insensitivity. Several studies have shown the cultivation of *ghd7* and *ghd7 dth7* genotypes in areas at high latitudes (>N53°), which have cool summers and a short growth season^17, 20^ (Fig. 8f). The *indica* cultivars containing the *hd1 dth8* double mutations also exhibit gradual daylength sensing and experience a shorter vegetative growth period, similar to rice cultivated in the late season at middle latitudes (Class 3), and still produce seeds (Fig. 8e–f and Supplementary Fig. 3).

At low latitudes (Classes 4 and 5), the genotypes with gradual daylength sensing also promote flowering and rapid life cycling for two-round planting (Fig. 8g–h and Supplementary Fig. 3). The *indica* cultivars carrying the *hd1* or *dth8* mutations with the fine-tuned critical daylength threshold (13 h) also constantly activate *Hd3a* expression at low latitudes (maximum daylength <13.5 h) in *indica* cultivars (Fig. 8g– h and Supplementary Fig. 18). These *indica* cultivars carrying the *hd1* or *dth8* mutations also adapt to low latitudes to promote flowering and a rapid life cycle for two-round planting (Fig. 8g–h and Supplementary Fig. 3), in addition to one-round planting at mid-latitudes (Fig. 8a, c and Supplementary Fig. 3). Therefore, the data confirm that fine-tuning response to critical daylength is an effective strategy to enhance crop latitude adaptation.

### Two Strategies for Accelerating Crop Latitude Adaptation Selection using DEAS

Recently, many models have shown that global warming has caused crop planting zones to shift northwards in the past few decades^1–6, 51, 52^. Future increases in temperature may open up new agriculturally suitable areas at high latitudes, and crops at middle and low latitudes will benefit from an extension in the growth period with temperature increases^1, 2^. Therefore, the selection of new crop varieties with accelerated latitude adaptation might be critical for yield stability in future crop migration. DEAS only requires about one month to establish the link between the daylength-sensing mechanism and latitude adaptability of crops, based on expression data for the florigen gene. Therefore, given the long duration and high economic cost of field trials to measure adaptability, DEAS offers a rapid and efficient method with which to accelerate adaptation selection for future food security and sustainable agriculture.

We suggest two strategies for currently incorporating DEAS into breeding programs. In crops such as rice, for which many photoperiod genes have been cloned, CRISPR/Cas9 genome editing technology can be used for for molecular breeding^53, 54^. The combination of DEAS and high-throughput sequencing can enable the daylength-sensing process and haplotypes of all photoperiod genes of the reference varieties at a given latitude to be obtained. Based on this information, the CRISPR/Cas9-mediated targeted knockout, knock-in or replacement of photoperiod genes can improve the adaptation of breeding material to a given latitude (Supplementary Fig. 23a).

Although no evidence demonstrates that potato *StCDF1*, soybean *Tof11, Tof12 and J*, maize *ZmCCT9*, *ZmCCT10* and *ZCN8* regulate daylength sensing to effect latitude adaptation in these crops, DEAS successfully detected daylength-sensing processes in maize and soybean (Fig. 7 and Supplementary Fig. 22), which shows that our approach can be extended to other crops. For new crops from which functional photoperiod genes have not been cloned, or for some crops (soybean and maize) from which some functional photoperiod genes have been cloned, DEAS can also play a predictive role by detecting the expression of the florigen gene to accelerate adaptation selection. Firstly, DEAS can be used to analyze reference varieties from different latitudes and to characterize the daylength-sensing features of these varieties at different latitudes. Secondly, DEAS can be used to classify breeding materials and populations according reference varieties for subsequent breeding (Supplementary Fig. 23b). Via this strategy, a defined breeding resource can be generated to accelerate adaptation selection under climate change in the future.

## Methods

### Generation of Genetic Material

Plants with the genotypes *Hd1 DTH8 Ghd7*, *Hd1 dth8 Ghd7, hd1 DTH8 Ghd7* and *hd1 dth8 Ghd7* were generated from the F_2_ generation of a cross between Dongjin (DJ) and *hd1 dth8*^22^. CRISPR/Cas9^55^ was used to generate *Hd1 DTH8 ghd7* and *hd1 dth8 ghd7* mutants in the DJ background. Specific target sites were designed using an online toolbox (http://crispr.dbcls.jp/)^56^. For *Hd1 DTH8 ghd7*, two unique 24-bp sequences, 5′-GGCATCCCCTGGCACGCACTCGGC -3′ and 5′-GCCGTATTGGATTGATACTCACGA -3′, were inserted into U3-gRNA and U6a-gRNA, respectively. These U3-gRNA and U6a-gRNA constructs were then cloned into the pYLCRISPR/Cas9-Mtmono vector and transformed into the Dongjin background by *Agrobacterium tumefaciens*-mediated transformation (*Agrobacterium* strain EHA105). For the *hd1 dth8 ghd7* mutants, three guide RNAs, respectively targeting *Hd1* (5′-GTTGGAGCGGCTGCCCATGGGCG -3′), Ghd7 (5′-GCCGATCGGCGCCCCGGCGCCAC -3′) and *DTH8* (5′-GGCAGCGCCGGGTATGTCGTCTA -3′) were cloned into U6b-gRNA, U6a-gRNA, and U3-gRNA. Subsequent steps followed a process identical to that used to generate *Hd1 DTH8 ghd7*. The NIL*^Hd1 DTH8 Ghd7^*, NIL*^hd1 DTH8 Ghd7^*, NIL*^Hd1 dth8 Ghd7^* and NIL*^hd1 dth8 Ghd7^* genotypes correspond to previously reported *DDHH*, *DDhh*, *ddHH* and *ddhh*, respectively^22^. NIL*^hd1 DTH8 ghd7^* as a pure line was derived from an earlier-flowering plant (contain D62B *ghd7* allele) from the BC5F2 population between parental cultivars D62B (indica) and 93-11 (indica, recurrent parent). Flowering time showed a segregation ratio of 3:1 in F2 population of NIL*^hd1 DTH8 ghd7^* and 93-11 (Supplementary Fig. 25a). Thus, the flowering time phenotype in the F2 population was regulated by a single gene. Further, we analyzed 768 plants from that population showing the recessive phenotype and found co-segregation between these phenotypes and two markers: RM5438 and AK61, which are located in *Ghd7* (Supplementary Fig. 25b). Comparative sequencing of the *Ghd7* alleles in 93-11, D62B, Zhenshan97^20^ and NIL*^hd1 DTH8 ghd7^* revealed that the Ghd7-containing genomic fragment was completely deleted in D62B, Zhenshan97 and NIL*^hd1 DTH8 ghd7^*. Further, the sequencing analysis showed NIL*^hd1 DTH8 ghd7^* harbors a 93-11 *DTH7* allele. NIL*^Hd1 dth8 Ghd7^* is also 93-11 background^22^. 93-11 harbors a non-functional *DTH7* allele^15^. NIL*^Hd1 DTH8^* ^gh*d7*^, NIL*^Hd1 dth8 ghd7^* and NIL*^hd1 dth8 ghd7^* were pure lines derived from the F_2_ generation of a cross between NIL*^hd1 DTH8 ghd7^* and NIL*^Hd1 dth8 Ghd7^*. Thus, all NILs contain homozygous non-functional *DTH7* alleles (Supplementary Fig. 25c). Primers are listed in Supplementary Table 1.

### Daylength-sensing-based Environment Adaptation Simulator (DEAS)

In DEAS Step1, seedlings from the DJ background and *indica* hybrid rice seedlings were grown in one of seven growth chambers at 28°C for 28 days; each growth chamber provided a different daylength (16 h, 14.5 h, 14 h, 13.5 h, 13 h, 12 h, or 10 h). At 29 days, seedlings in growth chambers with daylengths of 14.5 h, 14 h, 13.5 h, 13 h, and 12 h were divided into three groups per growth chamber. Two groups of seedlings were transferred to two neighboring growth chambers with increased or decreased daylengths, and the third group continued to grow in the original chamber. Seedlings in growth chambers with 16 h and 10 h daylengths were divided into two groups. One group of seedlings was transferred to the neighboring growth chamber and the other group continued to grow in the original growth chambers. After growth chamber transfer, all seedlings were exposed to the new daylength for 5 days and were then harvested. On day 6, all seedlings were harvested 3 h after dawn (ZT3) (Fig. 3a). Transcriptional data for *Ghd7*, *Ehd1*, *Hd3a*, and *RFT1* seedlings were then collected. When the expression difference of *Hd3a* or *RFT1* between two adjacent treatments in decreasing daylength was greater than ten-fold, we concluded that this expression pattern represented critical daylength sensing (Fig. 3c). When *Hd3a* or *RFT1* expression changed gradually with the change in daylength and the difference between adjacent treatments was less than ten-fold, we categorized this expression pattern as representing gradual daylength sensing (Fig. 3d). In DEAS Step2, *Hd3a* and *RFT1* expression data in increasing decreasing daylengths from DEAS Step1 can infer the dynamic of *Hd3a* and *RFT1* expression that corresponds to natural daylength changes at a specific latitude (Fig. 4a). The dynamics of *Hd3a* and *RFT1* expression at different latitudes can accurately describe the environmental adaptability of rice with different daylength-sensing processes on the map (Fig. 4a). The light in the growth chambers was supplied by LEDs (Sanan Sino-Science) and the LED spectrum is shown in Supplementary Fig. 26. A similar procedure was used for soybean and maize, but *Hd3a* and *RFT1* in rice were replaced by *GmFT2a* and *ZCN8*, respectively, in soybean and maize.

### Enrichment Analysis

We used a hypergeometric test to analyze the enrichment of different genotypes. *p*-values were calculated using the following formula:

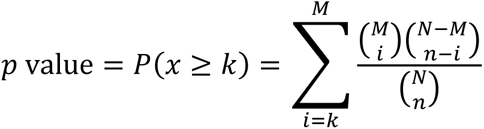

For Fig. 1b, *N* represents the number of all possible combinations of sterile lines and restorer lines, *n* represents the number of different genotypes in all possible combinations of sterile lines and restorer lines, *M* represents the number of *indica* hybrid rice varieties and *k* represents the number of different genotypes within the *indica* hybrid rice varieties.

For Supplementary Fig. 3, *N* represents the number of *indica* hybrid rice varieties, *n* represents the number of different genotypes in these varieties, *M* represents the number of *indica* hybrid rice varieties in different ecological classes and *k* represents the number of varieties with different genotypes in different ecological classes.

For Supplementary Figs. 5, 7 and 8, *N* represents the number of *indica* hybrid rice varieties in combined ecological classes, *n* represents the number of varieties with different genotypes in combined ecological classes, *M* represents the number of varieties that were planted in different ecological classes and *k* represents the number of varieties with different genotypes that were planted in different ecological classes.

### Simulated Ecological Classes for Daylength-Sensing Mechanisms

Seedlings from the DJ background were grown in five growth chambers at 28°C until flowering; each growth chamber provided a dynamic daylength to simulate a given ecological class. All seedlings were harvested 3 h after dawn (ZT3) on the given days (Fig. 5). Next, *Hd3a* or *RFT1* transcriptional data were collected from the seedlings. The light in the growth chambers was provided by LEDs (Sanan Sino-Science); the LED spectrum is shown in Supplementary Fig. 26.

### Categorical Manner for Evaluating DEAS Inference Accuracy

In DEAS Step2, *Hd3a* and *RFT1* expression data in increasing and decreasing daylengths from DEAS Step1 can infer a single dynamic of *Hd3a* and *RFT1* expression that corresponds to natural daylength changes at one specific latitude (Fig. 4a). We divided the expression of florigen genes into three intervals in the inferred *Hd3a* and *RFT1* expression profiles. When the *Hd3a* expression was <10 × 10^−4^ and the *RFT1* expression was <10 × 10^−5^, we defined them as being inactivated. When 10 × 10^−4^ < *Hd3a* expression < 100 × 10^−4^ and 10 × 10^−5^ < *RFT1* expression < 100 × 10^−5^, we defined them as being weakly activated. When *Hd3a* expression was >100 × 10^−4^ and *RFT1* expression was >100 × 10^−5^, we defined them as being highly activated (Fig. 4a). When we used our data for the accuracy evaluation, they did not require any preprocessing, but when using published data for the accuracy evaluation, the minimum value of the data was first converted to the order of 10^−4^ (*Hd3a* expression) or 10^−5^ (*RFT1* expression) by multiplying by 10^n^. Subsequently, the data were converted by multiplying by 10^n^. If the original or converted expression data for any given daylength were exactly within the inference interval, the inference was considered successful; otherwise, it was considered a failure. Finally, the percentage 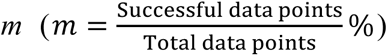 represented the DEAS inference accuracy in the categorical manner.

### Continuous Manner for Evaluating DEAS Inference Accuracy

To evaluate DEAS inference accuracy quantitatively, we generated a continuous manner. Firstly, at a given latitude, the expression value of *FT* orthologs at any time point in a year can be approximately inferred based on DEAS Step1. For example, using the Sun Day Length App developed by Sergey Vdovenko (http://www.lifewaresolutions.com/), we confirmed that the daylength in a paddy field at Tsukuba, Japan was 13.38 h on 19 August. This value was in the interval between 13.5 to 13 h; therefore, the expression value of *Hd3a* (Nipponbare background) in decreasing daylength (13.5 to 13 h) approximately represented *Hd3a* expression (Nipponbare background) on 19 August at this specific location. Furthermore, the maxima of the experimental data and inferred data were normalized to 1. Other data were converted by multiplying by a conversion coefficient 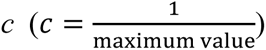. Next, using a linear regression model in R software, the coefficient of determination 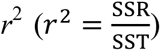 and *p*-value was calculated to evaluate the DEAS prediction accuracy (SST = total sum of squares; SSR = sum of squares due to regression).

For Fig. 6g–n, *Hd1 DTH8 Ghd7*, *Hd1 dth8 Ghd7*, *hd1 DTH8 Ghd7*, and *hd1 dth8 Ghd7* were grown in Chengdu, China under natural LDs (May to September). Leaves were harvested once a week at the same time (10:00) until the plants flowered.

### Fisher’s Exact Test

To detect whether temperature and other factors affect the DEAS prediction accuracy in the paddy field, Fisher’s exact test was used to calculate the probability of obtaining the observed data, and all data sets with more extreme deviations, under the null hypothesis that *m* _field_ and *m* _simulated classes_ are the same. If the *p*-value is <0.05, the result indicates that the null hypothesis can be rejected, i.e., that there is a significant difference in the DEAS inference accuracy in the categorical manner between simulated classes and the paddy field. If 0.05 *< p*-value *<*1, the *p*-value represents the probability that the DEAS inference is consistent with that of the simulated classes and the paddy field. The *p*-value was calculated using R software. The *m* _field_ and *m* _simulated classes_ were calculated in the categorical manner.

### Identification of Functional and Non-functional Alleles

All tested sterile and restorer lines were genotyped using the Illumina sequencing platform. From these sequencing data, the haplotypes of flowering-time genes in this study were collected^36^. We classified frameshift mutations and premature stop codons as non-functional alleles of *Hd1*, *DTH7*, *DTH8* and *Ghd7* and a crucial E105K mutation as non-functional *RFT1* allele; other haplotypes were classified as functional alleles. For partially sterile and restorer lines, whose alleles could not be determined by sequencing because of missing data, leaf tissue was frozen and DNA was extracted using a DNeasy Plant Mini kit (Qiagen). Gene fragments of *Hd1*, *DTH8*, *Ghd7*, and *DTH7* were amplified by PCR and sequenced with the primers listed in Supplementary Table 1. Sequencing results were analyzed by BLAST to ascertain the genotype of the *Hd1*, *DTH8*, *Ghd7* and *DTH7* alleles. The haplotype information for sterile and restorer lines can be obtained from MBKBASE (http://www.mbkbase.org).

### Data Collection

Information about *indica* hybrid rice varieties grown in East Asia, including days to heading and geographic distribution, was obtained from the China Rice Data Center (http://www.ricedata.cn/). Daylength data for Chinese provinces were collected using the Sun Day-Length App developed by Sergey Vdovenko (http://www.lifewaresolutions.com/).

### Analysis of Gene Expression

Total RNA was extracted from seedlings using the Eastep® Super Total RNA Extraction Kit (Promega), following the manufacturer’s instructions. cDNA was synthesized from 800 ng total RNA using GoScript^TM^ Reverse Transcription Mix using oligo(dT) (Promega). Real-time quantitative PCR was performed by the Taq-Man (rice) and SYBR Green (soybean and maize) PCR method on a CFX384^TM^ Real-Time PCR System (Bio-Rad) according to the manufacturer’s instructions. The primers used for PCR are listed in Supplementary Table 1.

### Yeast Two-Hybrid Assay

The full-length coding sequences (CDSs) of *DTH8*, *Ghd7*, and *Hd1* were amplified from *Nipponbare* cDNA and cloned into the *Eco*RI and *Xho*I sites of the pB42AD and pLexA vectors. Combinations of LexA BD- and AD- fusion plasmids were cotransformed into yeast strain EGY48 (Clontech) containing the reporter plasmid p8op-LacZ. Transformants were plated in darkness on uracil-, tryptophan-, and histidine-negative synthetic dropout medium containing X-gal (5-Bromo-4-chloro-3-indolyl-β-D-galactopyranoside) to visualize the development of blue color. The primers used are listed in Supplementary Table 1.

### Luciferase Complementation Imaging (LCI) Assay

Firefly LCI was performed in *Nicotiana benthamiana* leaves. The *DTH8* CDS sequence was ligated into the *Kpn*I/*Hind*III sites of the 35S:cLUC vector and the *Ghd7* CDS sequence was ligated into the *Kpn*I/*Sal*I sites of the 35S:cLUC and 35S:nLUC vectors. For the Hd1-nLUC construct, the *Hd1* CDS was ligated into the *Bam*HI/*Sal*I sites of the 35S:nLUC vector. *Agrobacterium* cultures (strain GV2260) were transformed with the nLUC- or cLUC-fused constructs and infiltrated into *N. benthamiana* leaves. After infiltration, the *N. benthamiana* plants were grown for 3 days, injected with luciferin and imaged using a Tanon 5200S Luminescent Imaging Workstation.

## ACKNOWLEDGMENTS

We thank the Yaoguang Liu lab for providing the pYLCRISPR/Cas9-MTmono vectors. We thank the breeder Lin Wang for providing the *indica* hybrid rice seeds. We thank Professor Xing Wang Deng, Professor Haiyang Wang and Professor Fanjiang Kong for reading and commenting on the manuscript. We thank Zengzheng Kui, Yanhui Wang, Weijuan Hu and Xin Liu for their technical assistance.

## Author Contributions

X.O. designed the research; L.Q., Q.W., X.W., G.Z., Z.S., J.H., H.W., W.T., Q.L., J.R., J.X., C.L., Y.L., S.L., R.H., X.C., C.Z., M.L., X.H., S.L. and X.O. performed the research; L.Q., P.Q., Z.C., Z.L., H.H., J.H., X.W., C.L. and X.O. analyzed the data; X.O. wrote the paper. The authors declare no conflicts of interest.

---

**Fig. S1.**
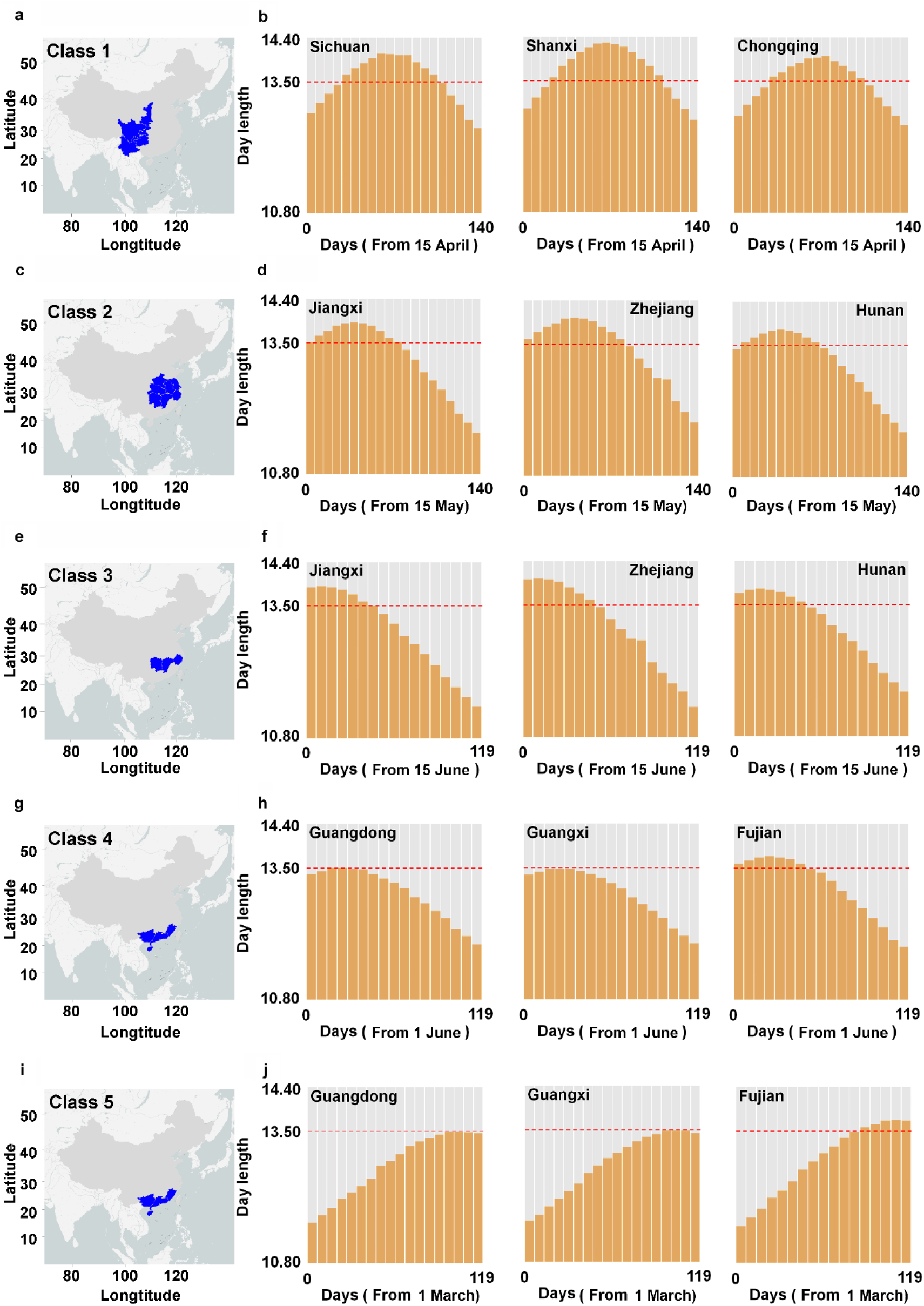
Five ecological classes of *indica* hybrid rice cultivation in East Asia. **a, c, e, g** and **i** Maps illustrating the five ecological classes. The blue areas on each map represent the geographic distribution of each class. **b, d, f, h** and **j** Photoperiod profiles at three collection sites within each ecological class. Yellow columns represent daylength and gray columns represent night length. Daylength data were collected every 7 days. The red dotted line indicates the critical daylength threshold. All maps were drawn using ArcGIS 10.3 software and modified using Photoshop CC2017.

**Fig. S2.**
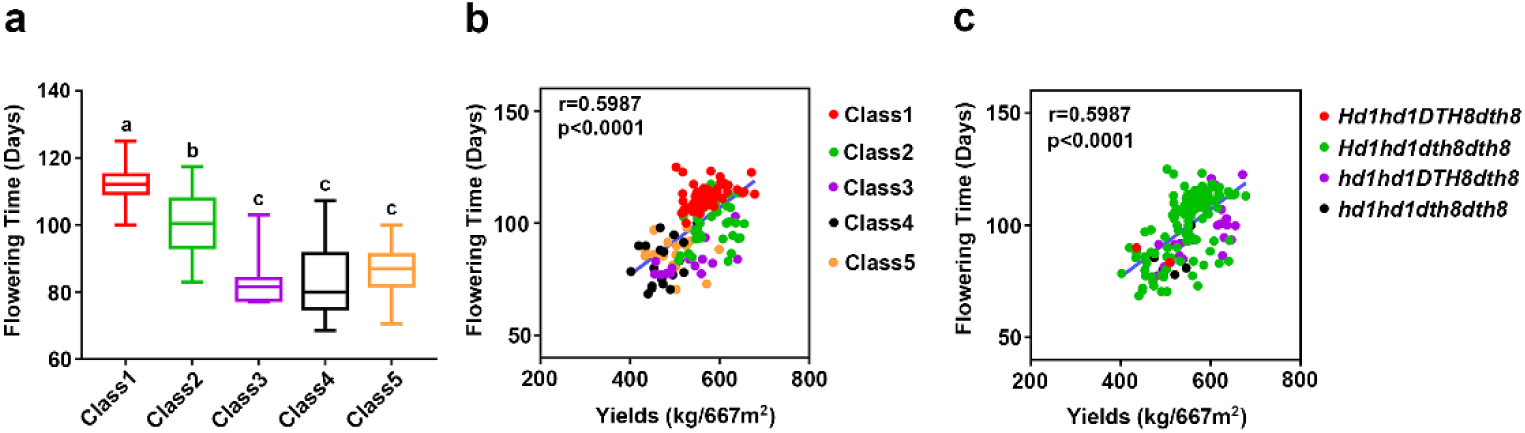
Relationship between flowering time and yield in *indica* hybrid rice varieties across ecological classes. **a** Flowering time in *indica* hybrid rice varieties across ecological classes. Data were analyzed using Duncan’s multiple range test (*P* < 0.05). Thin lines represent the flowering time of the same genotype across ecological classes. **b** The correlations between flowering time and yields in *indica* hybrid rice varieties across ecological classes. **c** The correlations between flowering time and yields in *indica* hybrid rice varieties across different genotypes.

**Fig. S3.**
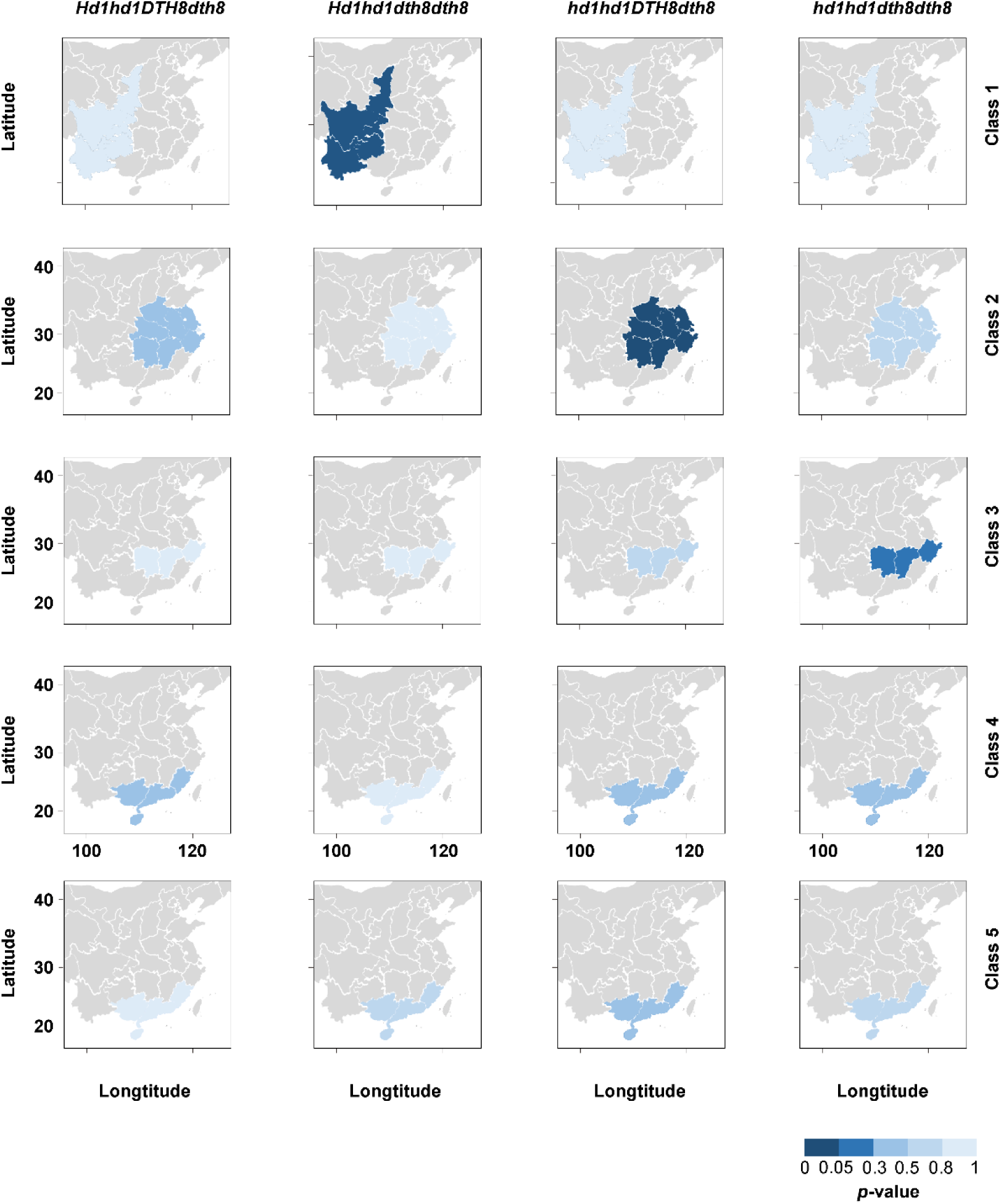
Enrichment of *Hd1* and *DTH8* allelic combinations in *indica* hybrid rice varieties across ecological classes. A heatmap representing enrichment of *Hd1* and *DTH8* allelic combinations in *indica* hybrid rice varieties within each ecological class. The *p*-values were calculated using hypergeometric analysis. All maps were drawn using ArcGIS 10.3 software and modified using Photoshop CC2017.

**Fig. S4.**
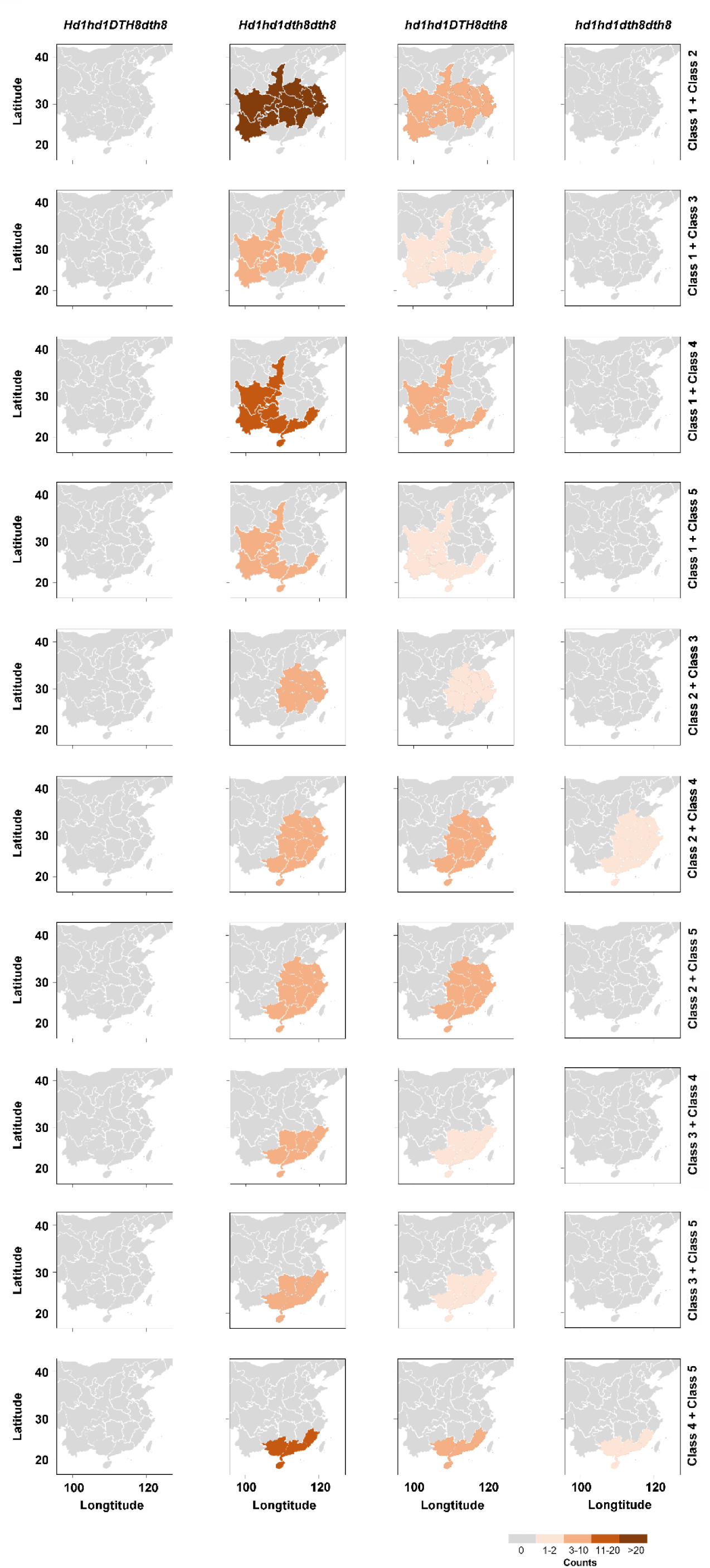
Heatmap to show the number of *Hd1* and *DTH8* allelic combinations in *indica* hybrid rice varieties in two combined ecological classes. All maps were drawn using ArcGIS 10.3 software and modified using Photoshop CC2017.

**Fig. S5.**
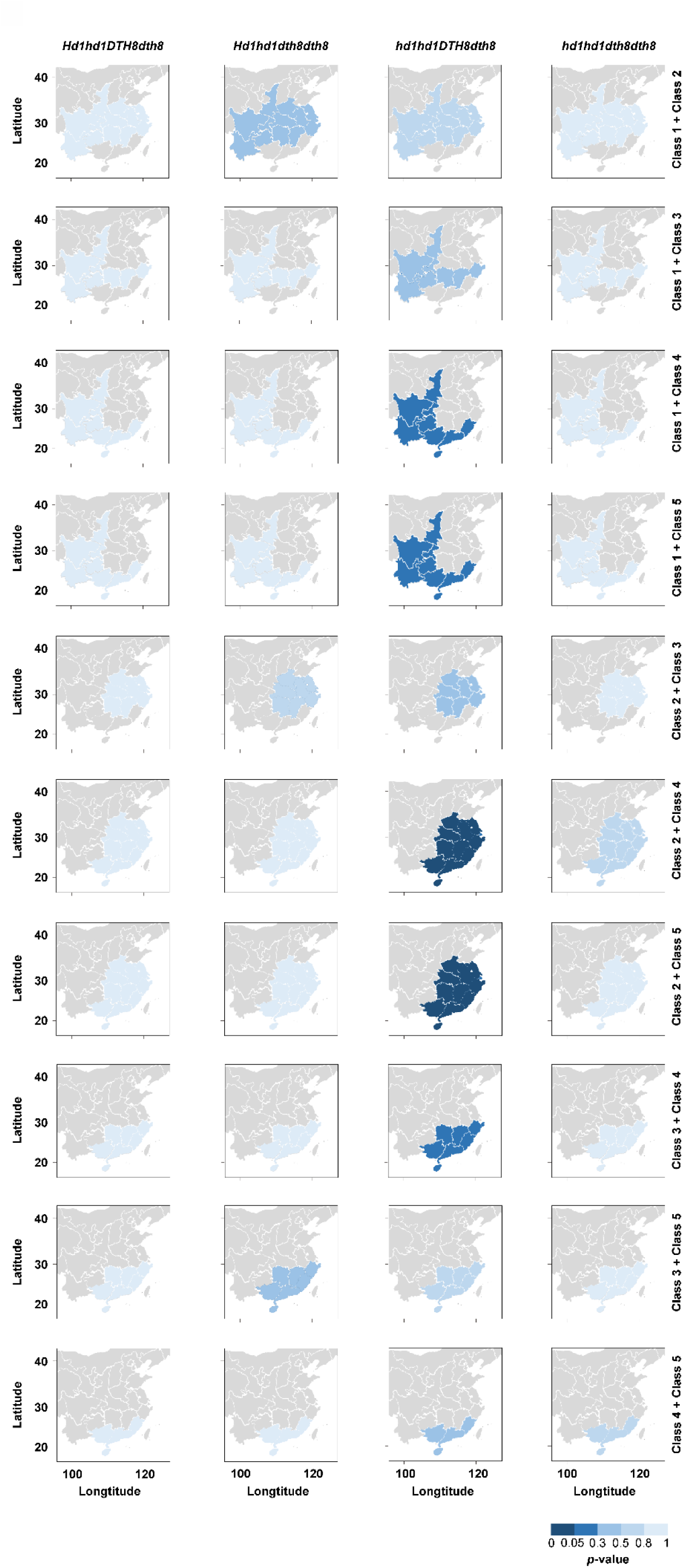
Heatmap to show the enrichment of *Hd1* and *DTH8* allelic combinations in *indica* hybrid rice varieties in two combined ecological classes. The *p*-values were calculated using hypergeometric analysis. All maps were drawn using ArcGIS 10.3 software and modified using Photoshop CC2017.

**Fig. S6.**
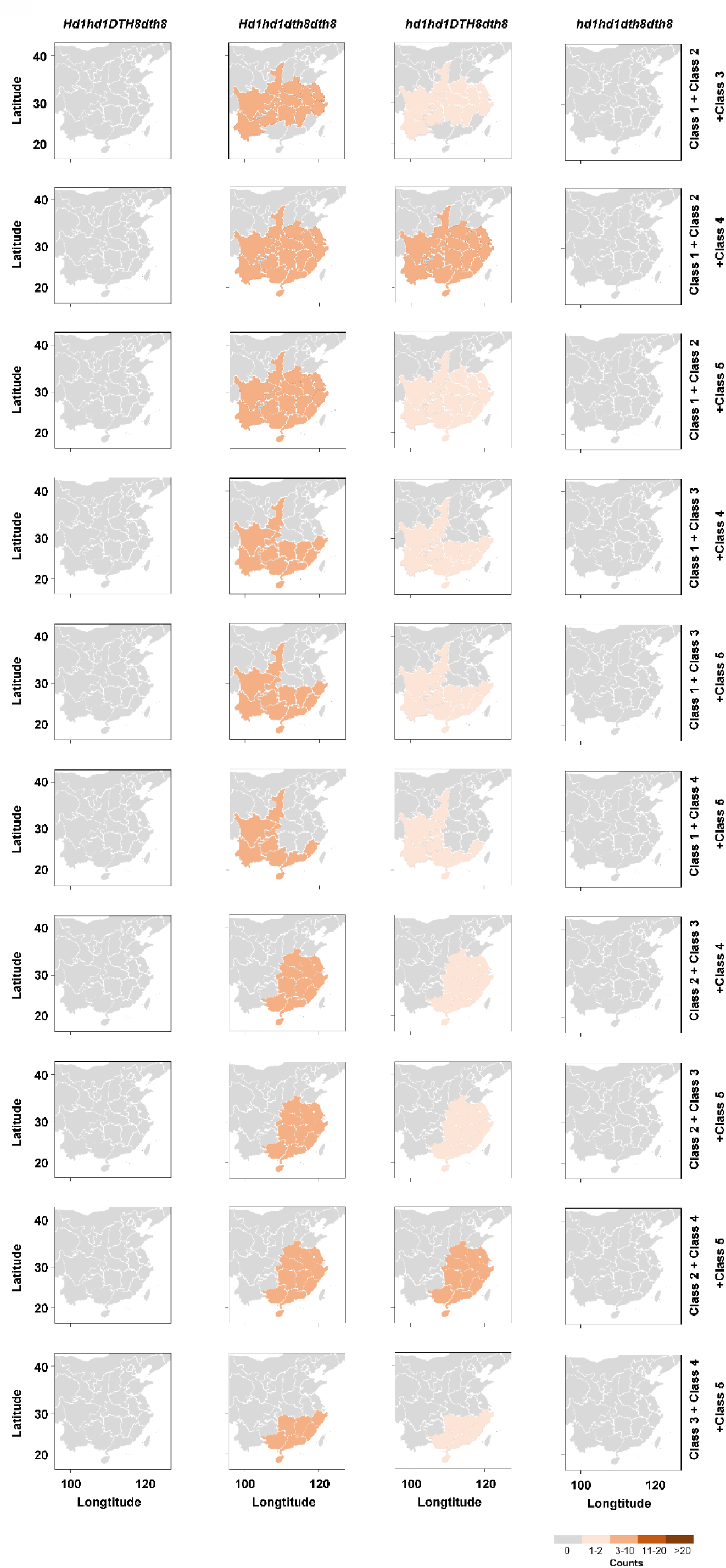
Heatmap showing the number of *Hd1* and *DTH8* allelic combinations in *indica* hybrid rice varieties in three combined ecological classes. All maps were drawn using ArcGIS 10.3 software and modified using Photoshop CC2017.

**Fig. S7.**
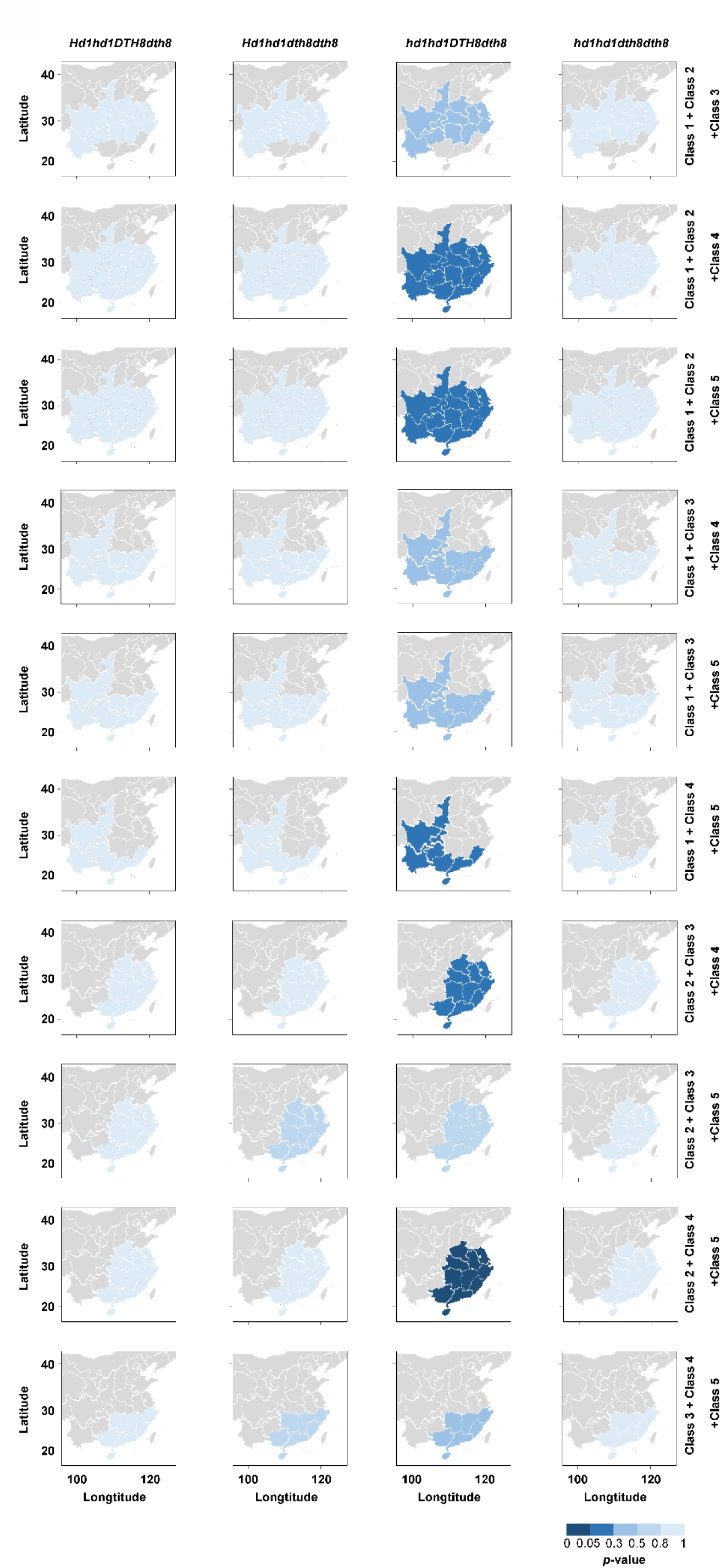
Heatmap showing the enrichment of *Hd1* and *DTH8* allelic combinations in *indica* hybrid rice varieties in three combined ecological classes. The *p*-values were calculated using hypergeometric analysis. All maps were drawn using ArcGIS 10.3 software and modified using Photoshop CC2017.

**Fig. S8.**
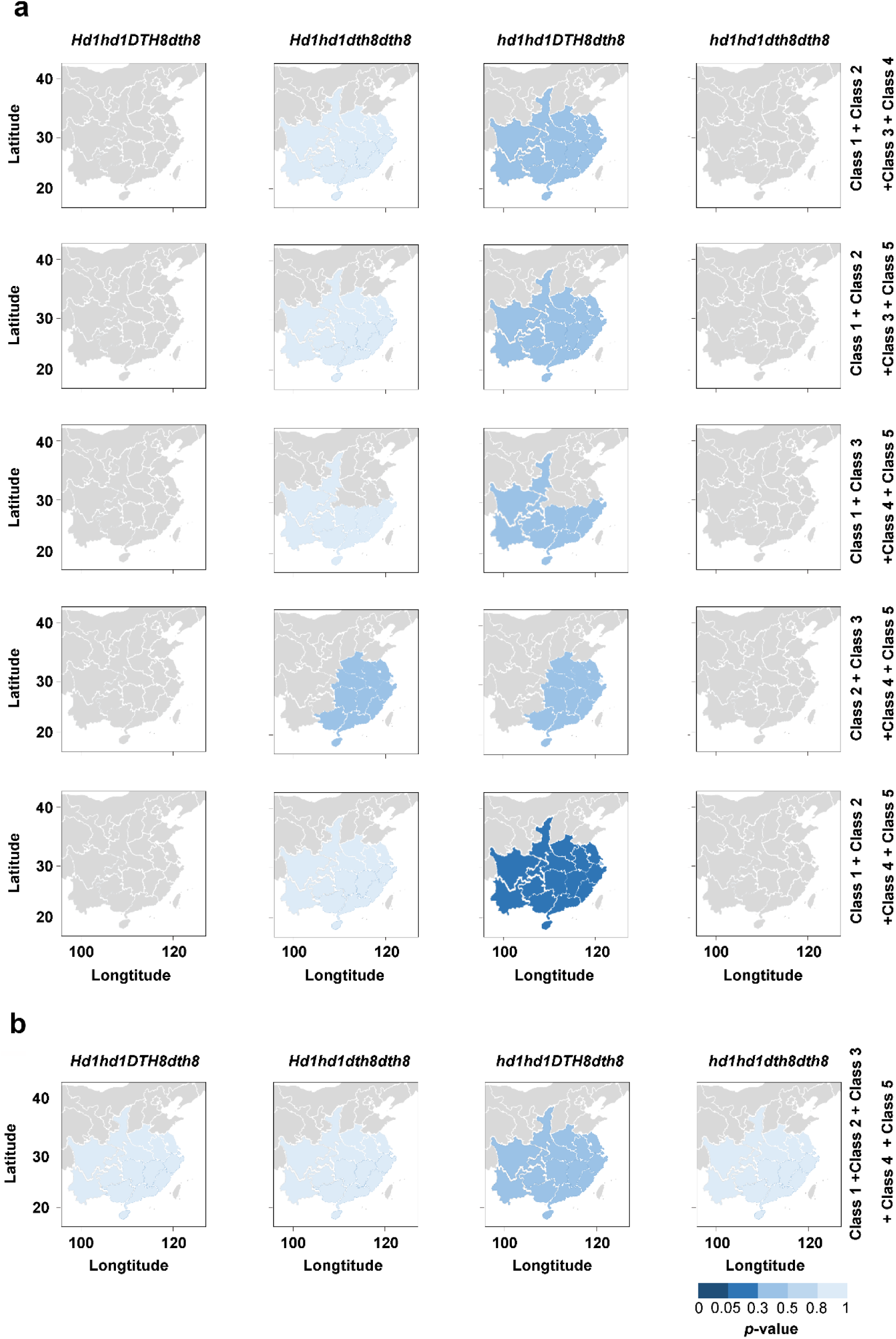
Enrichment of *Hd1* and *DTH8* allelic combinations in *indica* hybrid rice varieties within combined ecological classes. a A heatmap to represent the enrichment of *Hd1* and *DTH8* allelic combinations in *indica* hybrid rice varieties in four combined ecological classes. b A heatmap to represent the enrichment of *Hd1* and *DTH8* allelic combinations in *indica* hybrid rice varieties in five combined ecological classes. The *p*-values were calculated using hypergeometric analysis. All maps were drawn using ArcGIS 10.3 software and modified using Photoshop CC2017.

**Fig. S9.**
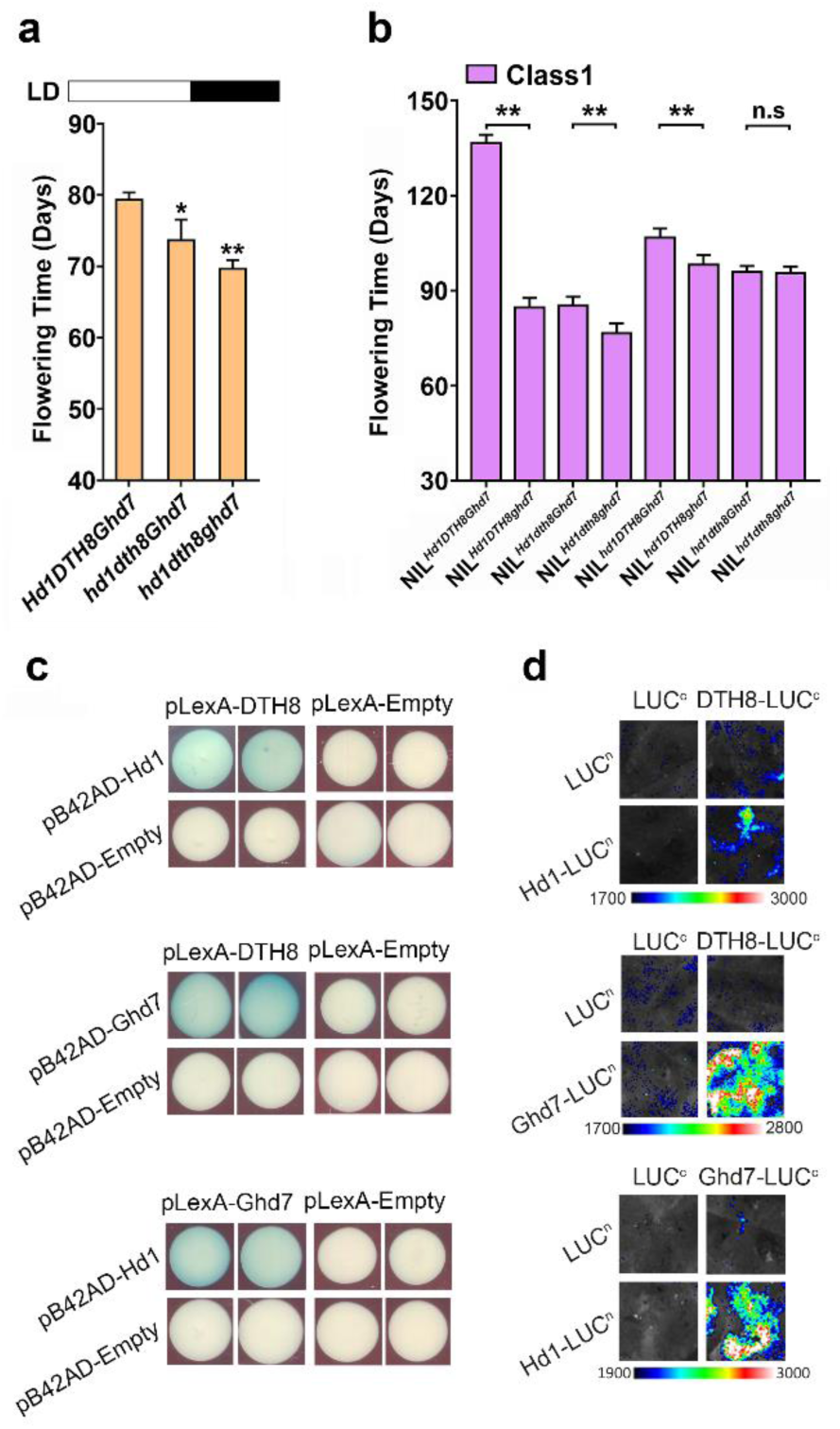
The Relationship between Ghd7 and DTH8-Hd1. **a** Flowering-time phenotypes of *Hd1 DTH8 Ghd7*, *hd1 dth8 Ghd7* and *hd1 dth8 ghd7*. Plants were grown under standard LD (14 h light:10 h dark at 28℃). Single and double asterisks above the graphs indicate statistically significant differences (*P* < 0.05 and *P* < 0.01, respectively). **b** Flowering-time phenotypes of NILs. Plants were grown in Class 1 (Chengdu, China) (Supplementary Fig. 1). Single and double asterisks above the graphs, indicate statistically significant differences (*P* < 0.05 and *P* < 0.01, respectively). **c** Yeast two-hybrid assay showing that Hd1, DTH8 and Ghd7 interact *in vitro*. **d** Luciferase complementation imaging (LCI) assay to confirm physical interactions among Hd1, DTH8, and Ghd7.

**Fig. S10.**
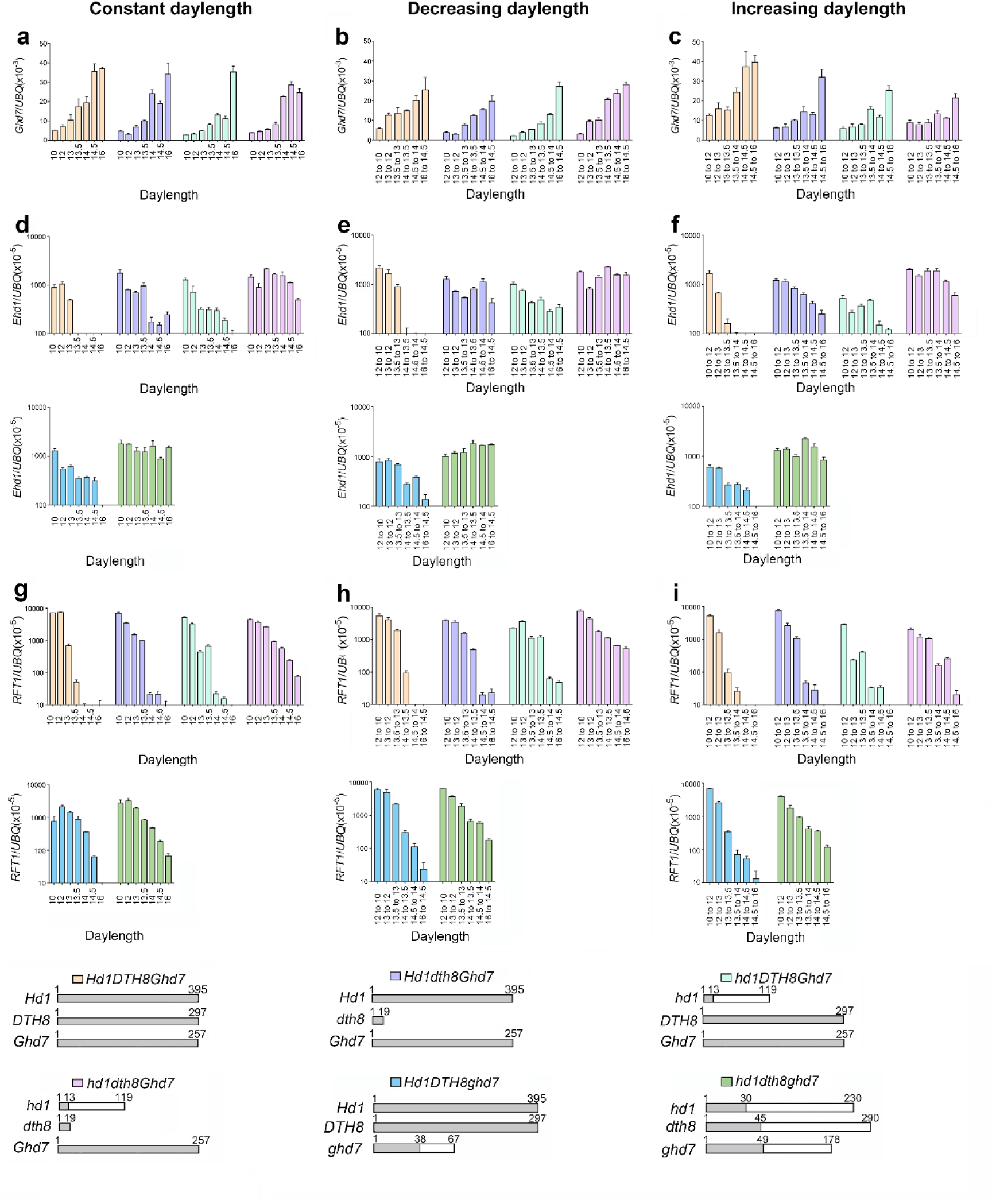
*Ghd7*, *Ehd1* and *RFT1* mRNA levels under various daylength conditions. Relative mRNA levels determined by RT-qPCR. Error bars represent the SD for three biological replicates. **a–c** *Ghd7* expression. **d–f** *Ehd1* expression. **g–i** *RFT1* expression. Gray rectangles show the lengths of the Hd1, DTH8 or Ghd7 proteins in numbers of amino acids, respectively. White rectangles represent proteins resulting from frameshift mutations in *Hd1*, *DTH8* or *Ghd7*. Small gray rectangles represent proteins resulting from a premature stop codon in *Hd1*, *DTH8* or *Ghd7*. DNA sequencing results are shown in Supplementary Fig. 24.

**Fig. S11.**
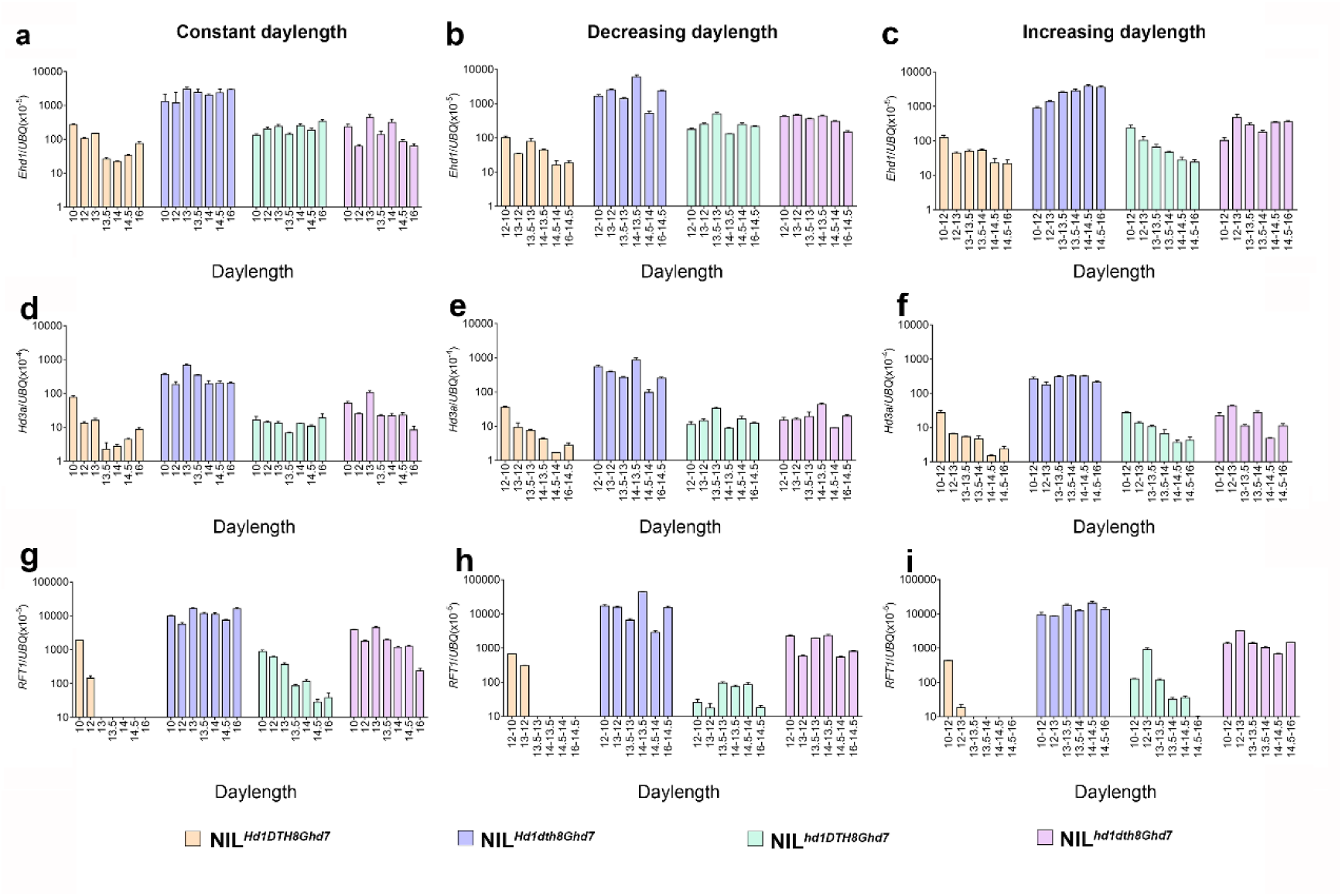
*Ehd1*, *Hd3a* and *RFT1* mRNA levels under various daylength conditions. Relative mRNA levels determined by RT-qPCR. Error bars represent the SD for three biological replicates. **a–c** *Ehd1* expression. **d–f** *Hd3a* expression. **g–i** *RFT1* expression.

**Fig. S12.**
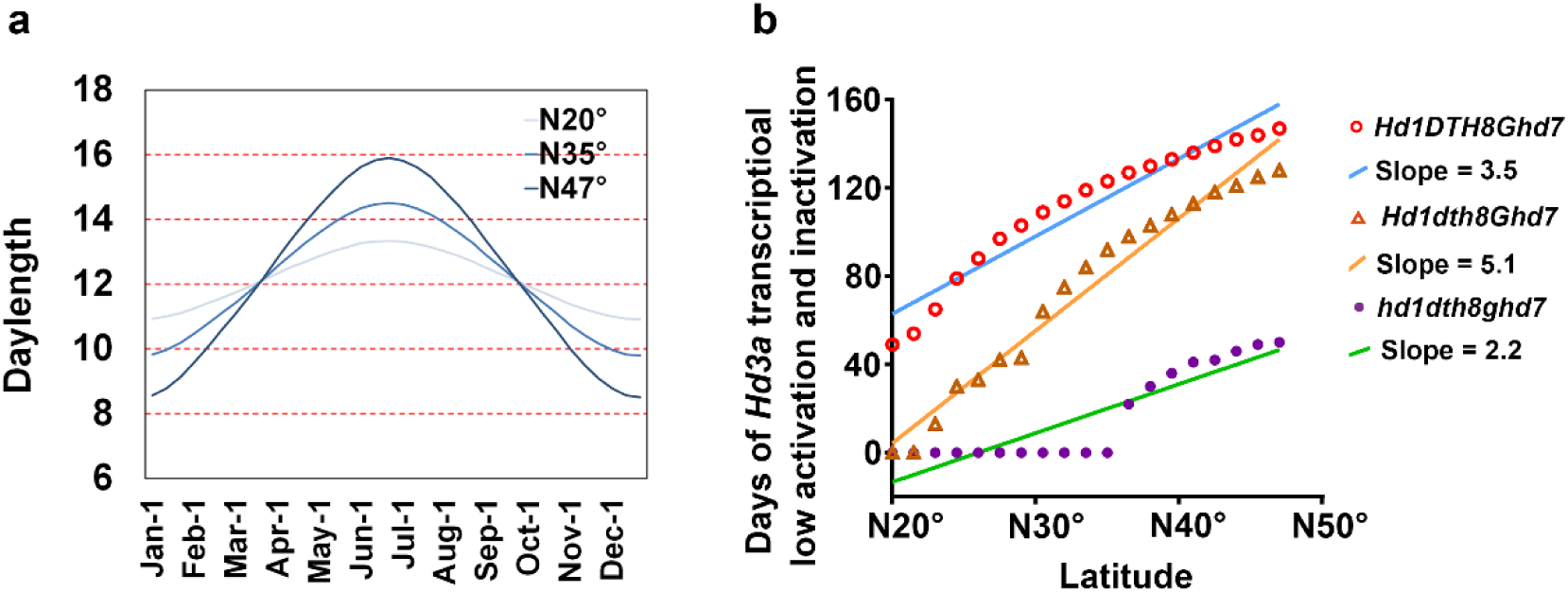
Relationship between daylength sensing and latitude in rice. **a** The Relationship between daylength and latitude. **b** The Relationship between the level of *Hd3a* transcription and latitude.

**Fig. S13.**
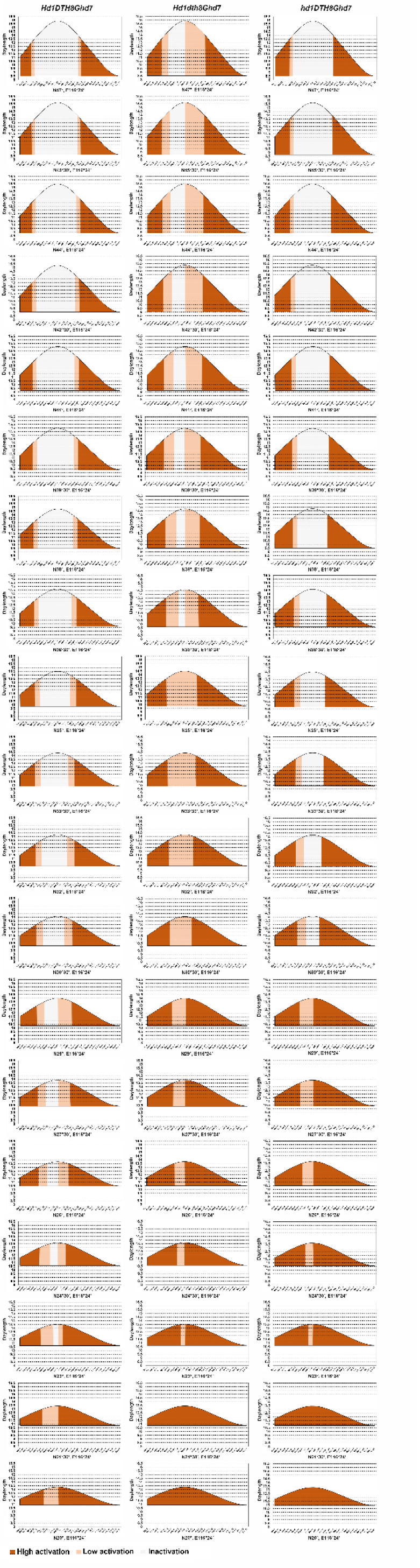
Dynamics of *Hd3a* transcription in response to different latitudes in genotypes with critical daylength sensing.

**Fig. S14.**
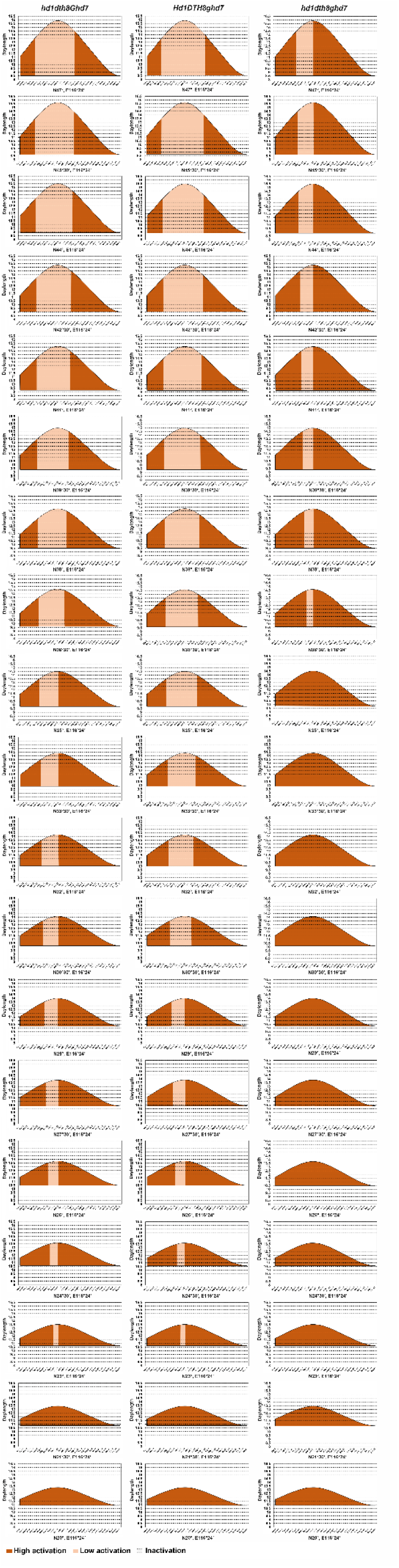
Dynamics of *Hd3a* transcription in response to different latitudes in genotypes with gradual daylength sensing.

**Fig. S15.**
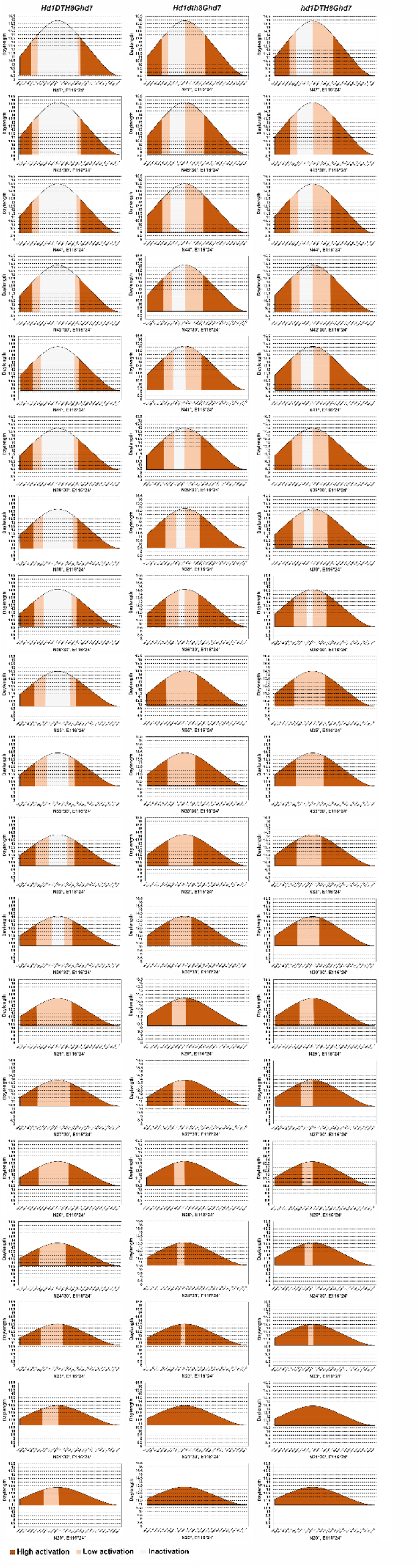
Dynamics of *RFT1* transcription in response to different latitudes in genotypes with critical daylength sensing.

**Fig. S16.**
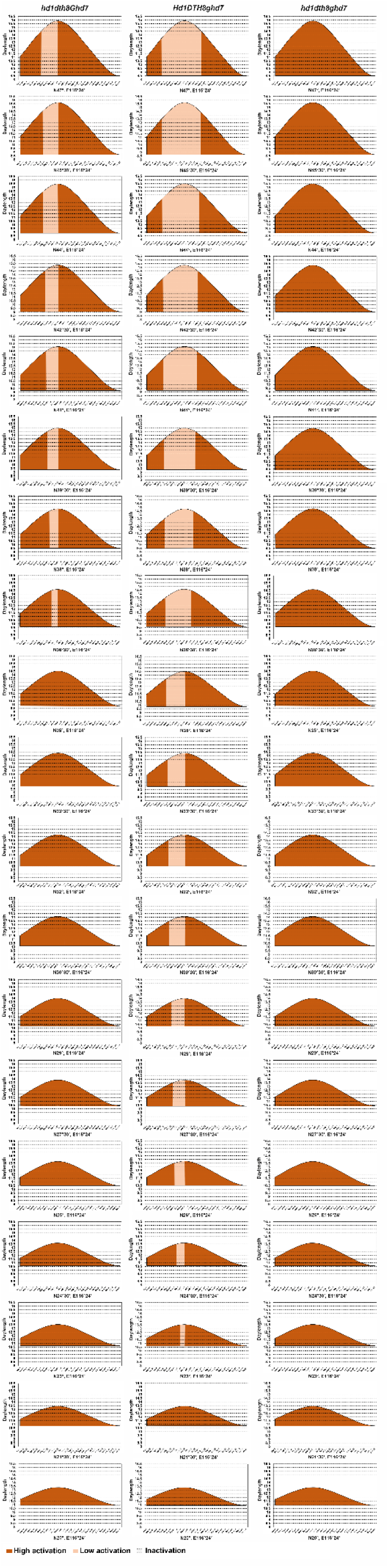
Dynamics of *RFT1* transcription in response to different latitudes in genotypes with gradual daylength sensing.

**Fig. S17.**
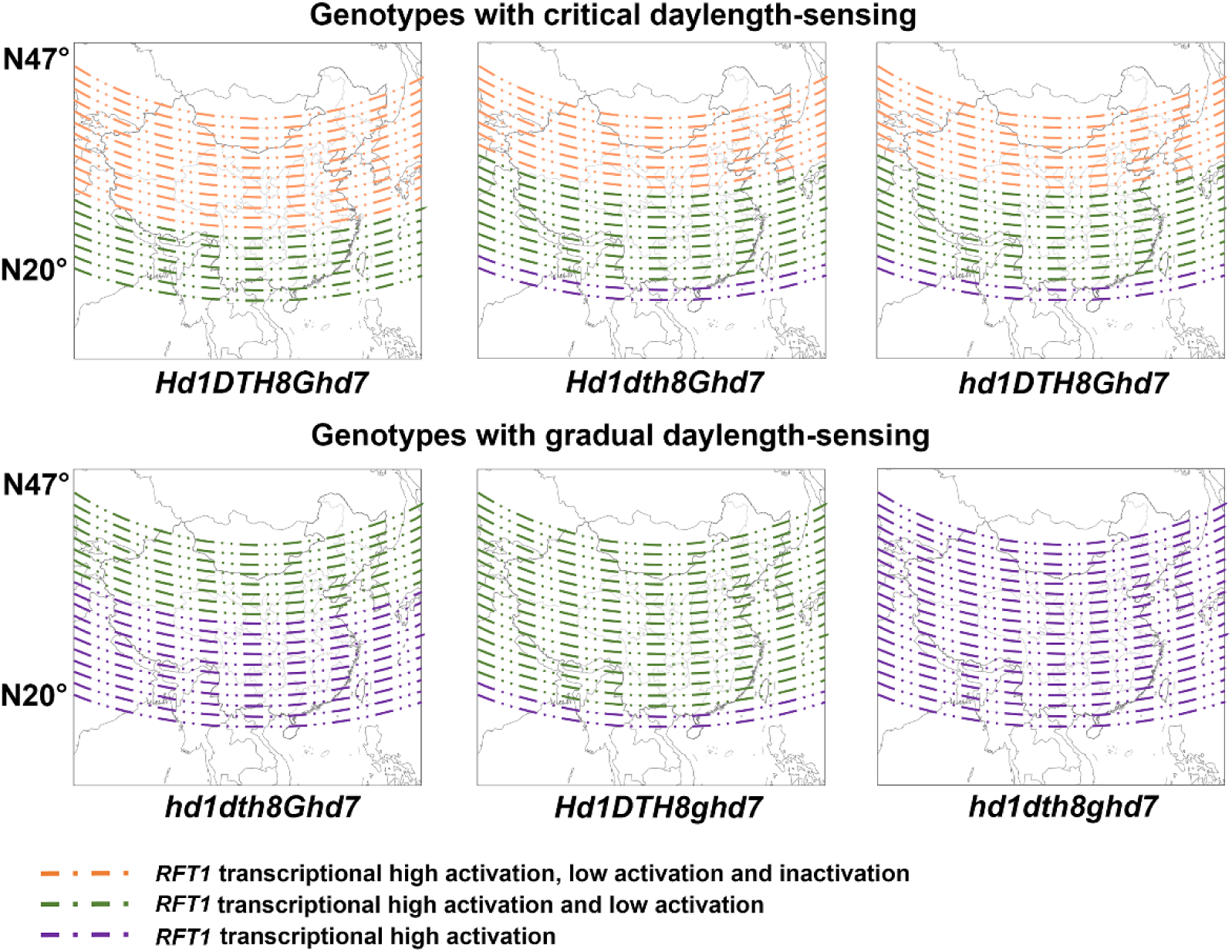
Maps illustrating different *RFT1* expression dynamics at different latitudes for different genotypes. All maps were drawn using ArcGIS 10.3 software and modified using Photoshop CC2017.

**Fig. S18.**
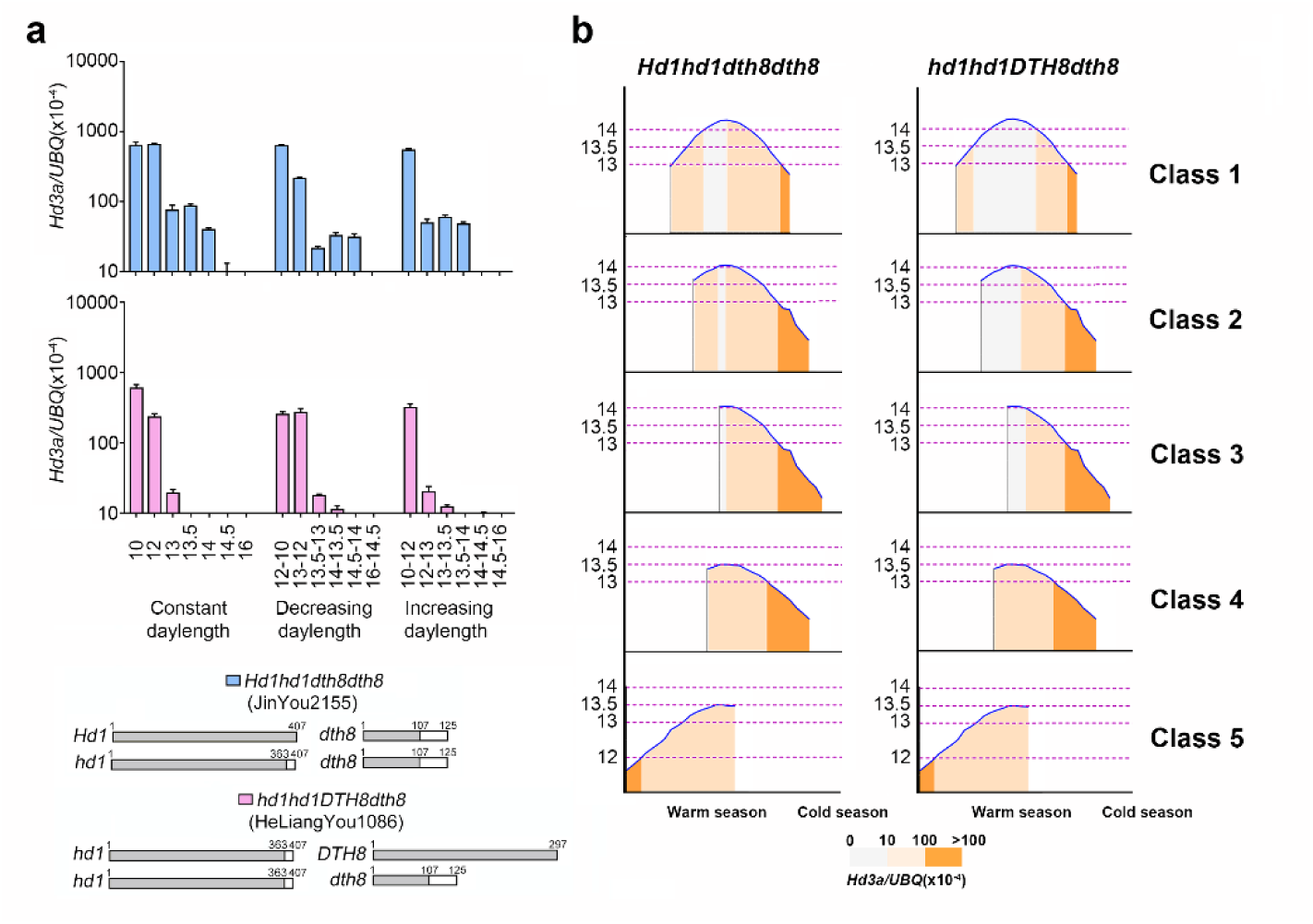
Daylength-sensing mechanisms in *indica* hybrid varieties within different ecological classes in East Asia. **a** *Hd3a* mRNA level under various daylength conditions. Relative mRNA levels determined by RT-qPCR. Error bars represent the SD for three biological replicates. Gray rectangles show the lengths of the Hd1 or DTH8 proteins in numbers of amino acids. White rectangles represent proteins resulting from frameshift mutations in *Hd1* or *DTH8*. **b** Daylength-sensing mechanisms in *Hd1hd1 dth8dth8* and *hd1hd1 DTH8dth8* within different ecological classes in East Asia. Blue lines indicate different daylengths. Different colored shading represents different *Hd3a* expression levels.

**Fig. S19.**
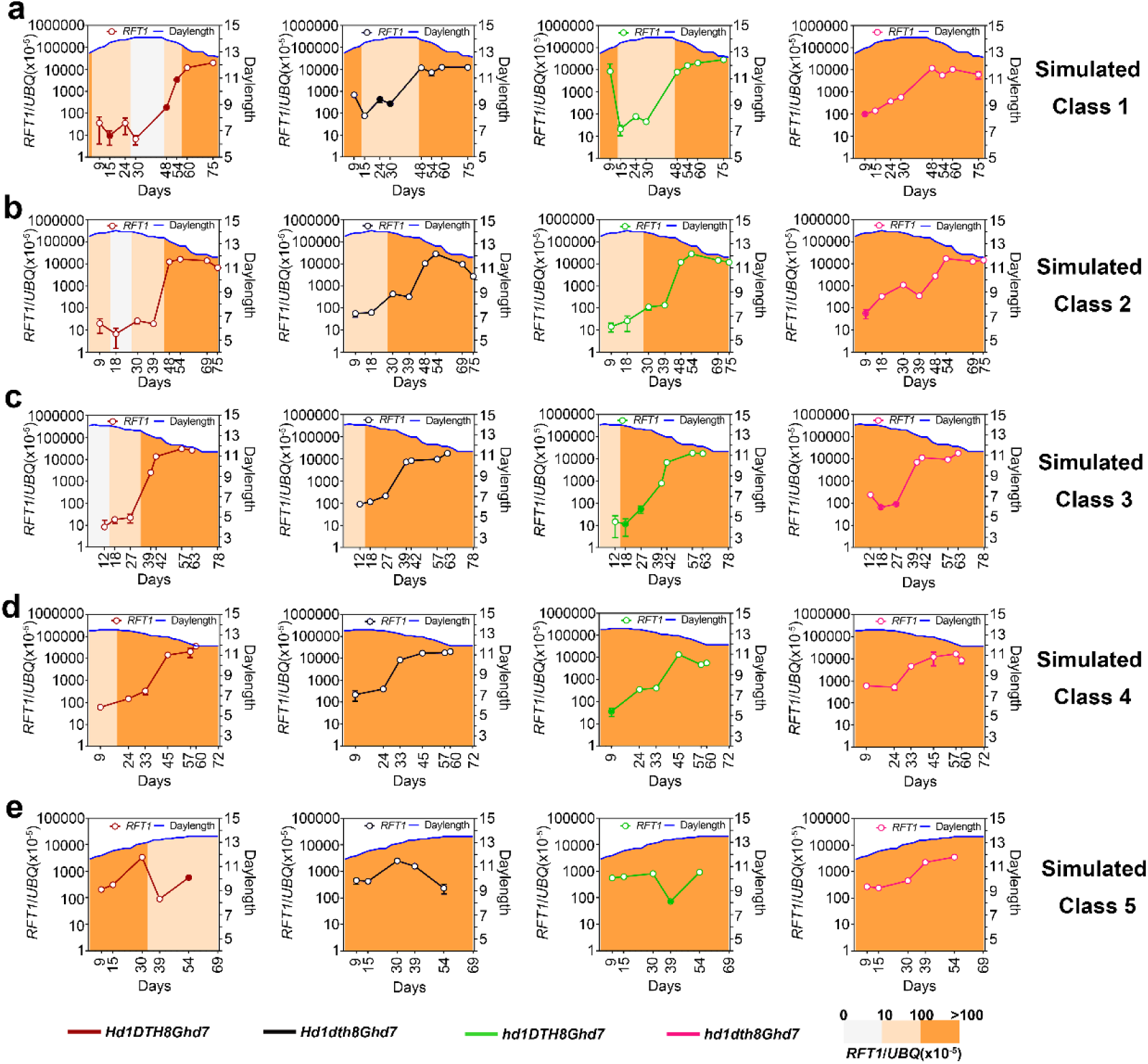
The categorical manner for evaluating DEAS inference accuracy using *RFT1* mRNA levels in simulated five ecological classes. The relative mRNA level determined by RT-qPCR. Error bars represent the SD for three biological replicates. Blue lines represent changing daylength in simulated five ecological classes. The four different colored lines represent the estimated value using DEAS. The filled circles and hollow circles indicate that the data were consistent or inconsistent, respectively, with the inference represented by three background colors.

**Fig. S20.**
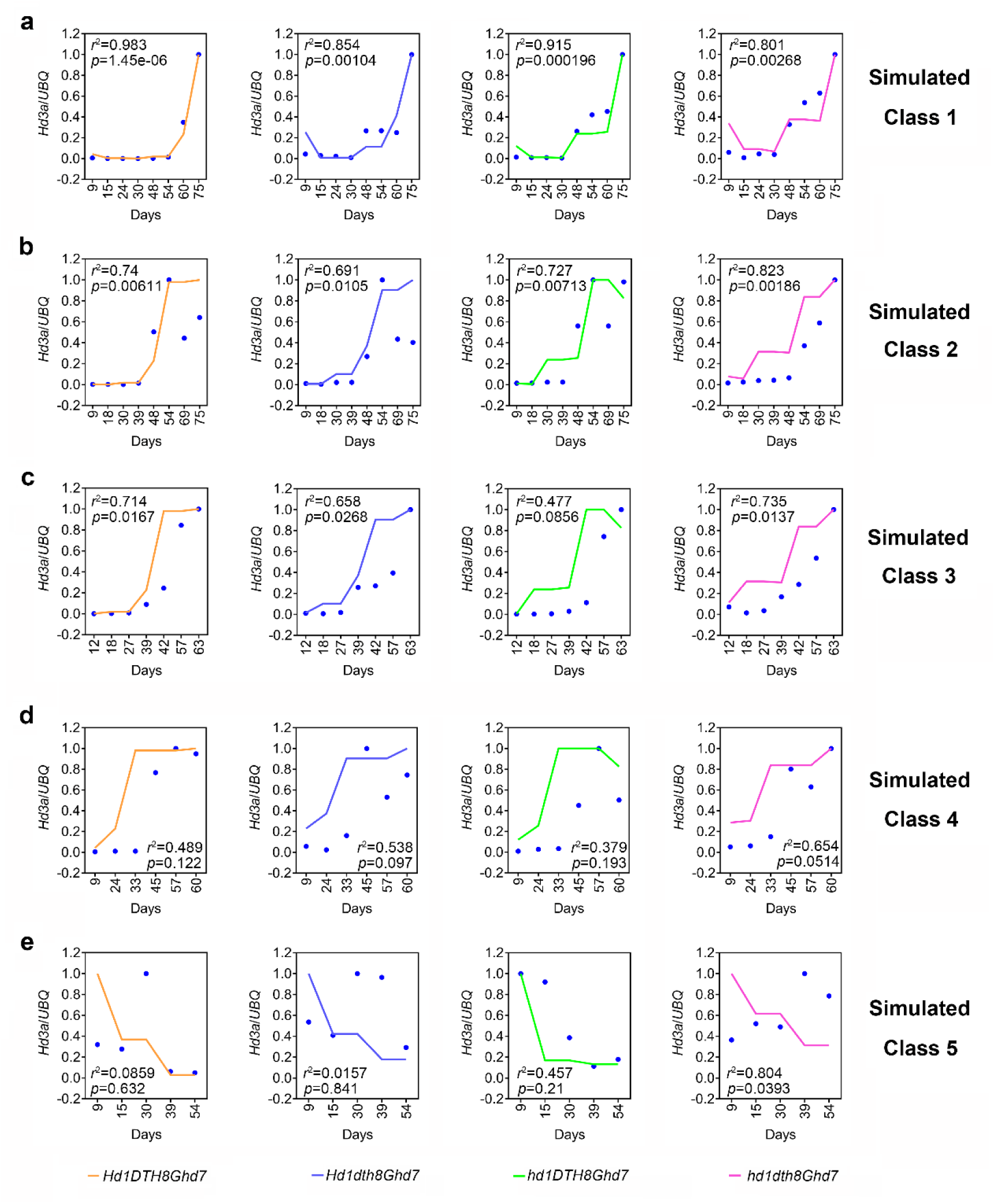
The continuous manner for evaluating DEAS inference accuracy using *Hd3a* mRNA levels in simulated five ecological classes. Blue points represent experimental data for *Hd3a* mRNA levels in five simulated ecological classes. Different colored lines represent the estimated value using DEAS. The coefficient of determination *r*^2^ and *p*-value can be calculated to evaluate DEAS inference accuracy.

**Fig. S21.**
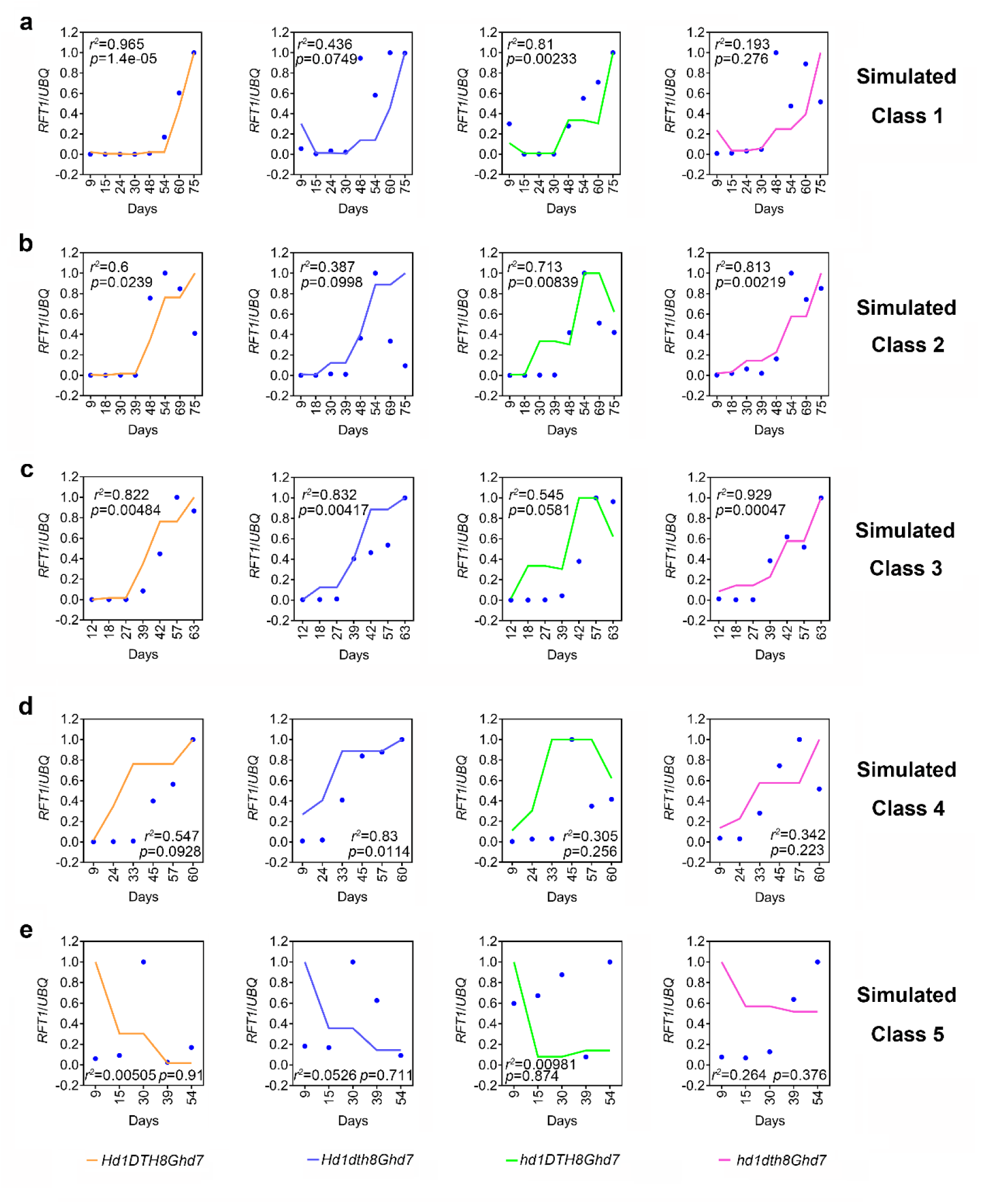
The continuous manner for evaluating DEAS inference accuracy using *RFT1* mRNA levels in five simulated ecological classes. Blue points represent experimental data for *RFT1* mRNA levels in five simulated ecological classes. Different colored lines represent the estimated value using DEAS. The coefficient of determination *r*^2^ and *p*-value can be calculated to evaluate DEAS inference accuracy.

**Fig. S22.**
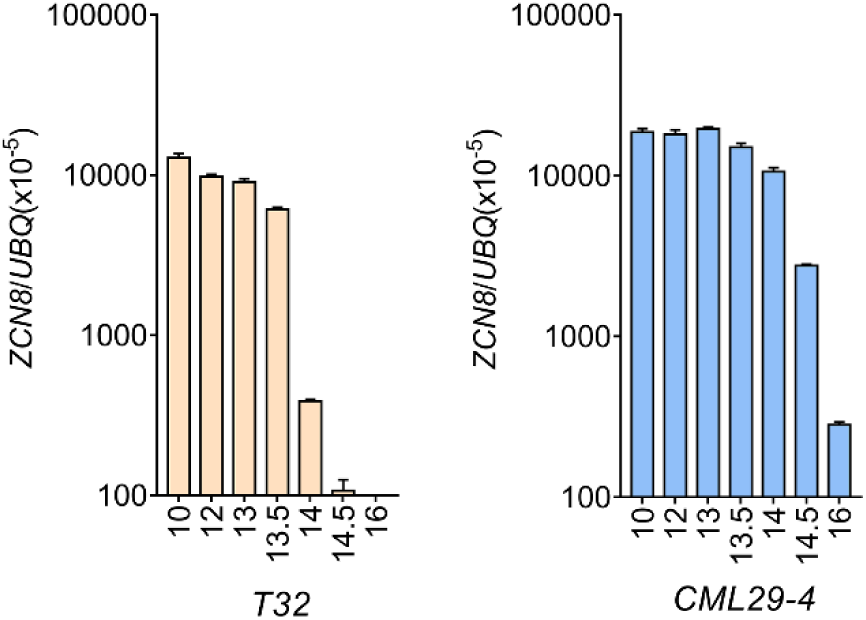
The application of DEAS to maize. The level of *ZCN8* mRNA under various daylength conditions in DEAS Step1. Relative mRNA levels determined by RT-qPCR. Error bars represent the SD for three biological replicates.

**Fig. S23.**
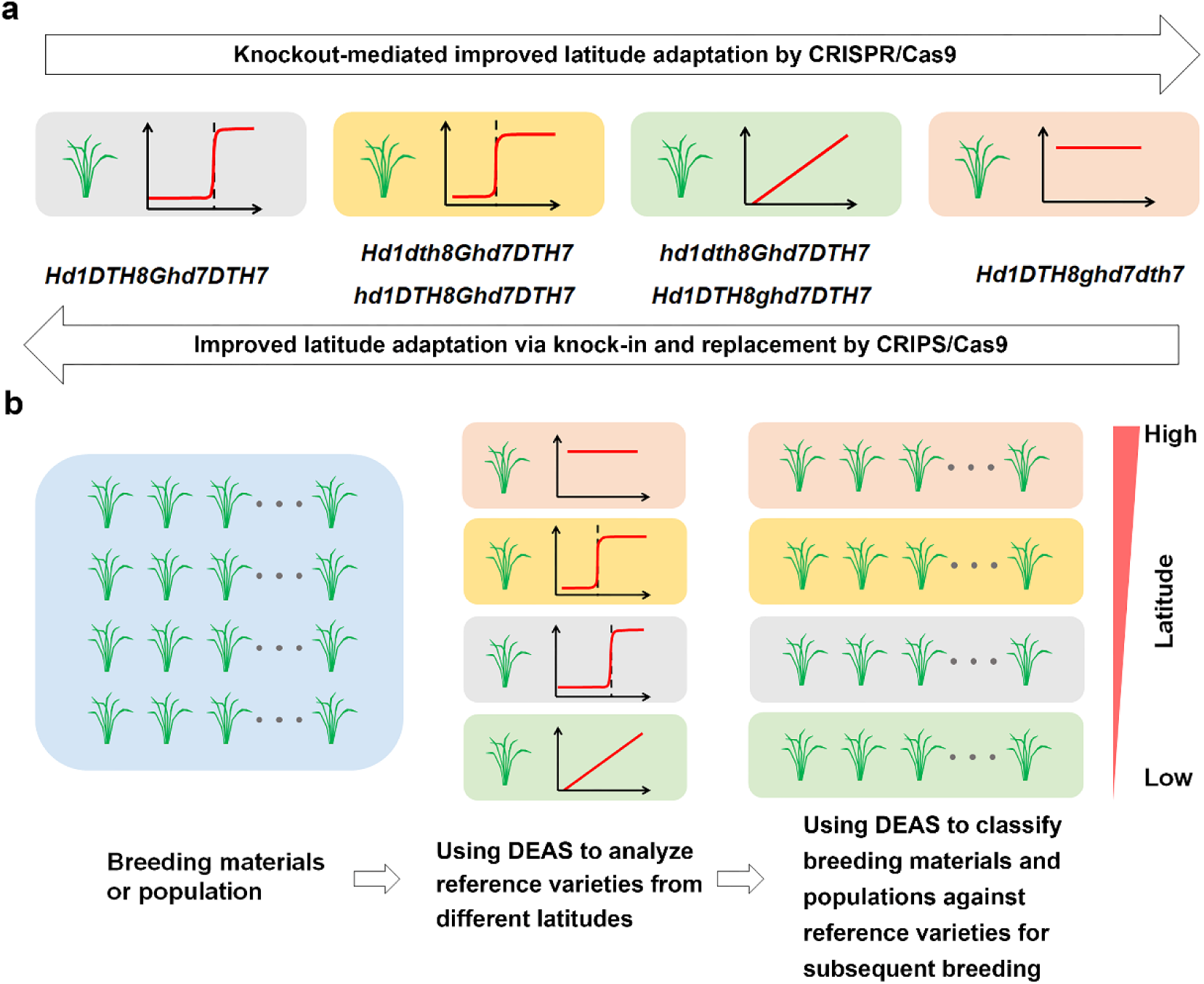
Two strategies for accelerating crop latitude adaptation selection using DEAS **a** CRISPR/Cas9-based crop latitude adaptation selection. CRISPR/Cas9-mediated targeted knockout, knock-in and replacement of photoperiod genes to improve the adaptation of breeding materials at a given latitude by changing daylength sensing. **b** Genetic population-based crop latitude adaptation selection. DEAS can be used to classify breeding genetic population against reference varieties for subsequent breeding.

**Fig. S24.**
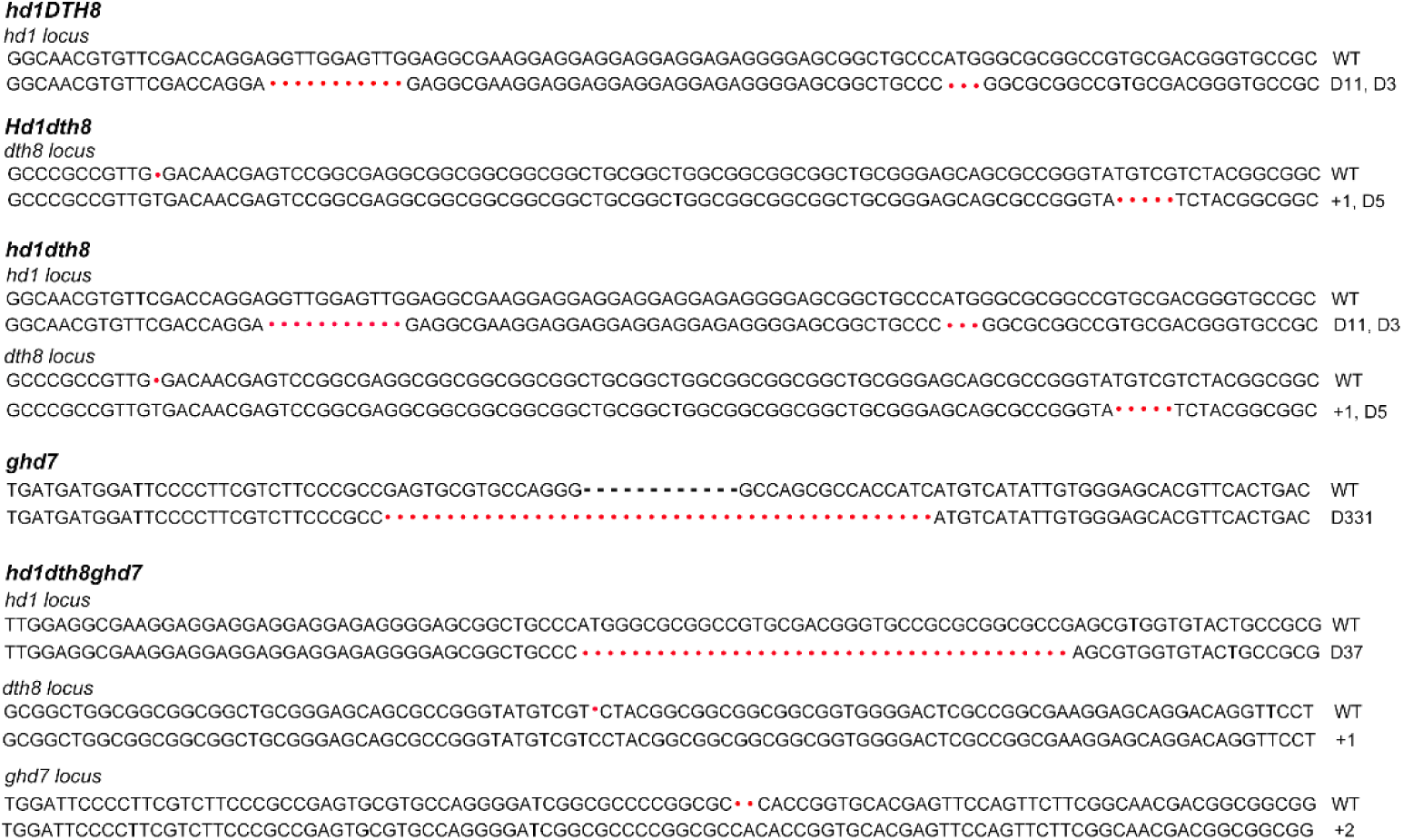
Sequence of different mutant alleles in the Dongjin background created using CRISPR/Cas9 technology.

**Fig. S25.**
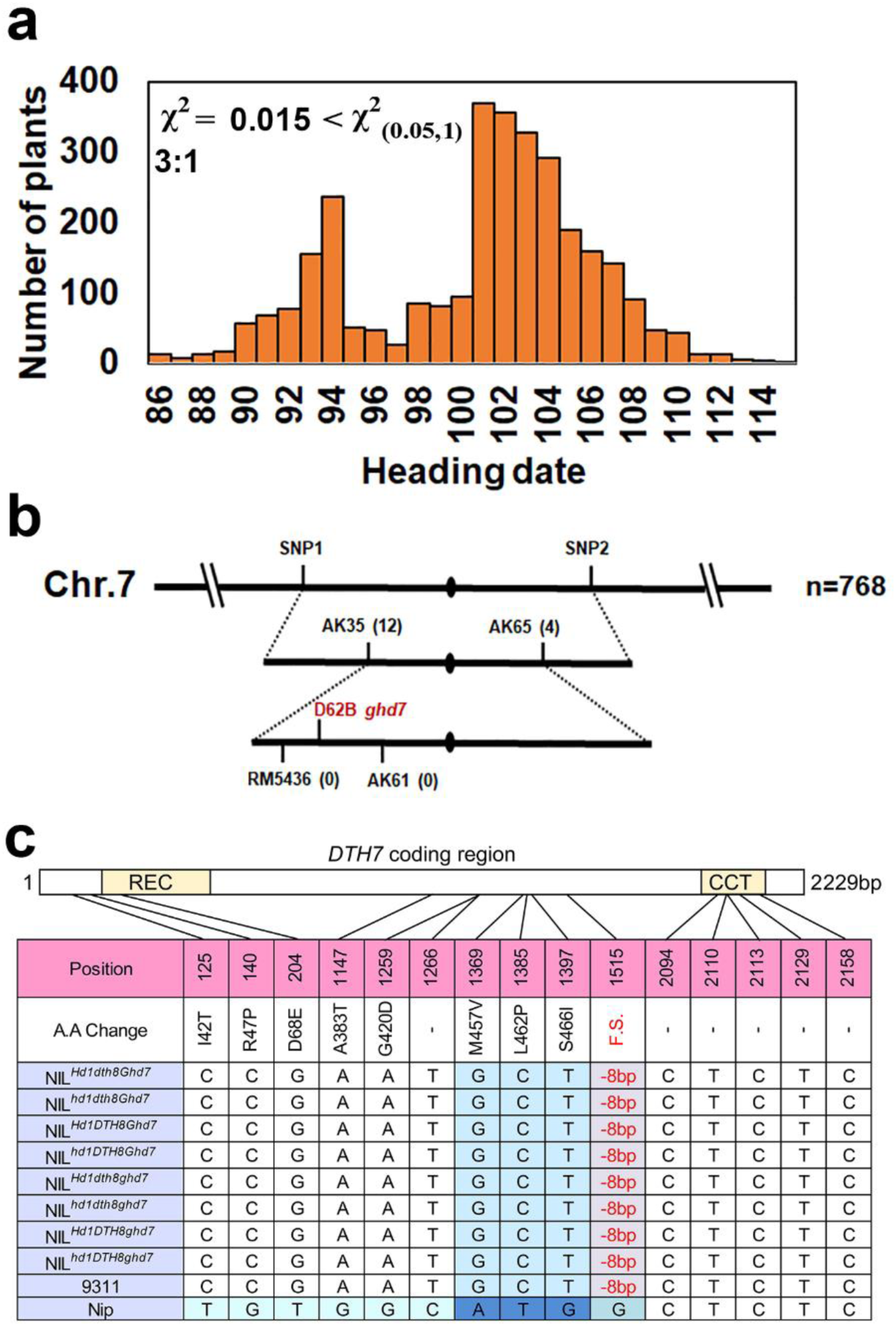
Genetic analysis of NILs. **a** Frequency distribution for flowering time in F2 population of NIL*^hd1 DTH8 ghd7^* and 93-11. Plants were grown under natural LD. Chi-square test results check the segregation ratio of flowering phenotypes in F2. **b** Fine mapping of *Ghd7*. **c** Different types of *DTH7* in NILs.

**Fig. S26.**
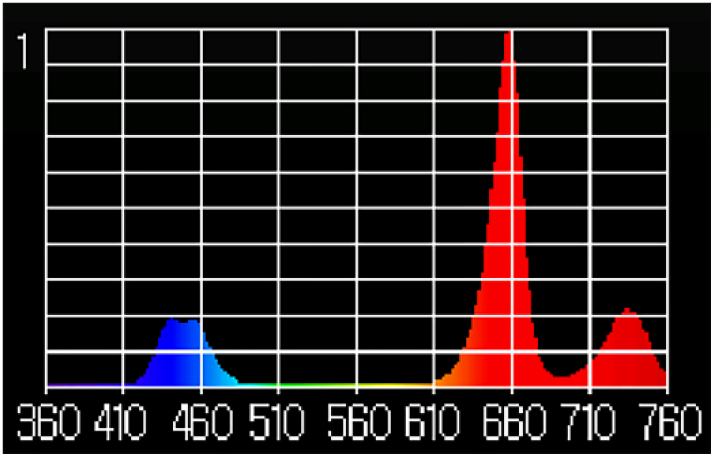
The light spectrum of LED (>450 µmol m^−2^ s^−1^) light in the growth chambers. Data were collected using a Hipot HR-350 light meter.

